# Taking fear back into the Marginal Value Theorem: the risk-MVT and optimal boldness

**DOI:** 10.1101/2023.10.31.564970

**Authors:** Vincent Calcagno, Frédéric Grognard, Frédéric M. Hamelin, Ludovic Mailleret

## Abstract

Foragers exploiting heterogeneous habitats must make strategic movement decisions in order to maximize fitness. Foraging theory has produced very general formalizations of the optimal patch-leaving decisions rational individuals should make. One is Charnov’s Marginal Value Theorem (MVT), which models the sequential visit of habitat patches and their spatial distribution. The MVT has a simple intuitive graphical interpretation in terms of gain functions and travel times. However, it considers only energy gains, and the effect of predation risk on the time allocation strategy is notoriously lacking. An important development that includes predation risk was Brown’s economic treatment of optimal patch leaving decisions, the basis of giving-up density (GUD) theory, often cited as an extension of the MVT. However, it is a more abstract result that does not have the specificities or graphical appeal of the MVT. Although both successful, the two theories are cited by distinct communities and are seldom connected in texbooks. Here we formally introduce the risk-MVT (rMVT), a generalization of the MVT that can incorporate most types of predation risks. We show that Brown’s GUD-theory is equivalent to a rMVT, but applies for one type of predation risk only. The rMVT retains the structure and graphical simplicity of the MVT, but implies a shift from residence time to expected dose of risk (micromort units, as used in decision analysis) as the domain over which rates of gain are computed and maximized. Applications of the rMVT show that different types of risk can yield opposite responses of optimal strategies to an increase in the risk level, and predict differential responses of behaviours observed in experimental versus natural conditions. The risk-MVT can also be used to predict the optimal level of risk taking, or “optimal boldness”, and suggests that individuals should generally be bolder in riskier habitats.

## INTRODUCTION

Optimal foraging theory has provided a rich framework to assess the behavior of individuals in the light of natural selection and adaptive evolution (Stephens and Krebs 1986; Begon et al. 1996; Mangel 2006). One very popular theory is the Marginal Value Theorem (MVT; Charnov 1976). The latter applies to a broad range of situations where individuals must decide between exploiting further their current patch of resources, or leaving the patch in search of another one (Calcagno 2018). The MVT models an explicit dynamics of sequential patch visits, whereby movement bouts and patch finding are a prerequisite to patch exploitation. The process of finding a new patch necessitates some amount of time, called the travel time. Patch exploitation also takes time, and the accumulation of gains in a patch is captured by the so-called gain function. This function can have various shapes, but generally describes diminishing returns, caused for instance by the gradual depletion of local resources (Calcagno et al. 2014). From a set of constraints imposed by habitat characteristics (spatial distribution and intrinsic values of resource patches), the MVT predicts the optimal time individuals should exploit a patch before leaving in search for another one (or residence time), and the fitness achieved in the habitat. The MVT thus connects local instantaneous decisions (when to leave a patch) to global properties of the habitat that can, at least in principle, be independently measured and manipulated. This coupling is best captured by the famous graphical solution of the homogeneous MVT, often presented in articles and textbooks (Parker and Stuart 1976; Stephens and Krebs 1986; Mangel 2006).

Although the MVT makes many simplifying assumptions, whose implications have been extensively discussed, its great simplicity probably underlies its sustained appeal. Nonetheless, a major limitation of the MVT is that it considers resource acquisition only, omitting the potential risks inherent to foraging, such as predation risk (Brown 1988; Calcagno 2018). This is quite problematic as for many taxa predation risk is a fundamental aspect of foraging: individuals can suffer attacks from predators and be perturbed by them in several ways, including death (Brown et al. 1999; Preisser et al. 2005) and a variety of less direct effects (Lima and Dill 1990). Foragers should sometimes disregard the most profitable habitat patches if these are too risky (Werner et al. 1983), but tolerate some level of risk in order to satisfy energy requirements (McNamara and Houston 1987; Verdolin 2006).

To address this shortcoming, another popular theory addressing patch-leaving decisions, usually called giving-up density (GUD) theory, was introduced by Brown (1988). Just like the MVT, GUD theory considers patchy habitats and predicts when individuals should leave a patch, based on marginal rates. Because of this similarity, the two theories are sometimes used more or less interchangeably regarding optimal patch leaving decisions. Still, they have significant differences. In particular, Brown’s theory characterizes an instantaneous decision at a particular point in time: at any time, the individual is seen as choosing between alternative activities, characterized by marginal gains or costs, with no need to specify what the activities are or imposing that some must be performed sequentially. It does not explicit how the different marginal rates arise from global habitat characteristics. In this respect, Brown’s theory is more general, and more abstract, than the MVT: it is a general description of the (local) mathematical properties of an optimal strategy. Regardless, the recognition that risk plays a role in shaping patch utilization patterns, and the possibility to include predation risk in foraging theory, was a major improvement brought forth by Brown’s (1988) theory, and the cause of its popularity.

Both theories have proved very successful, and both continue to serve as bases for further developments (Charnov and Parker 1995; Calcagno et al. 2013, 2014; Pacheco-Cobos et al. 2019; Arehart et al. 2023). As of May 2023, Charnov’s 1976 article had been cited 3,485 times and Brown’s 1988 article 1,031 times, according to the Scopus citation database (Fig. 1; see Supporting Information). This makes them some of the most cited papers in behavioral ecology and animal behavior ever, a remarkable achievement for theoretical pieces. Charnov’s 1976 publication is now regarded as iconic by its publishing journal (Rosenberg 2020). Both articles continue to be abundantly cited, at remarkably similar paces, their relative rate of citations quite stable over the past 20 years (Fig. 1a). The two theories thus appear to have reached some form of steady coexistence, but this coexistence does not imply much coalescence. Indeed, textbook treatments of the MVT often do not cover Brown’s developments (Begon et al. 1996; Westneat and Fox 2010), and reciprocally GUD-theory typically regards the MVT as an earlier step no longer to be considered in itself (Morris and Davidson 2000). In the literature, the two theories are generally not cited by the same articles or by the same authors (even though they are cited in similar journals; see Fig. S1). The relative overlap of their citing articles or citing authors, as quantified by Sorensen index, never exceeded a low 15% and are on a downward tendency (Fig. 1b), suggesting some divide in the community.

**Figure 1.**
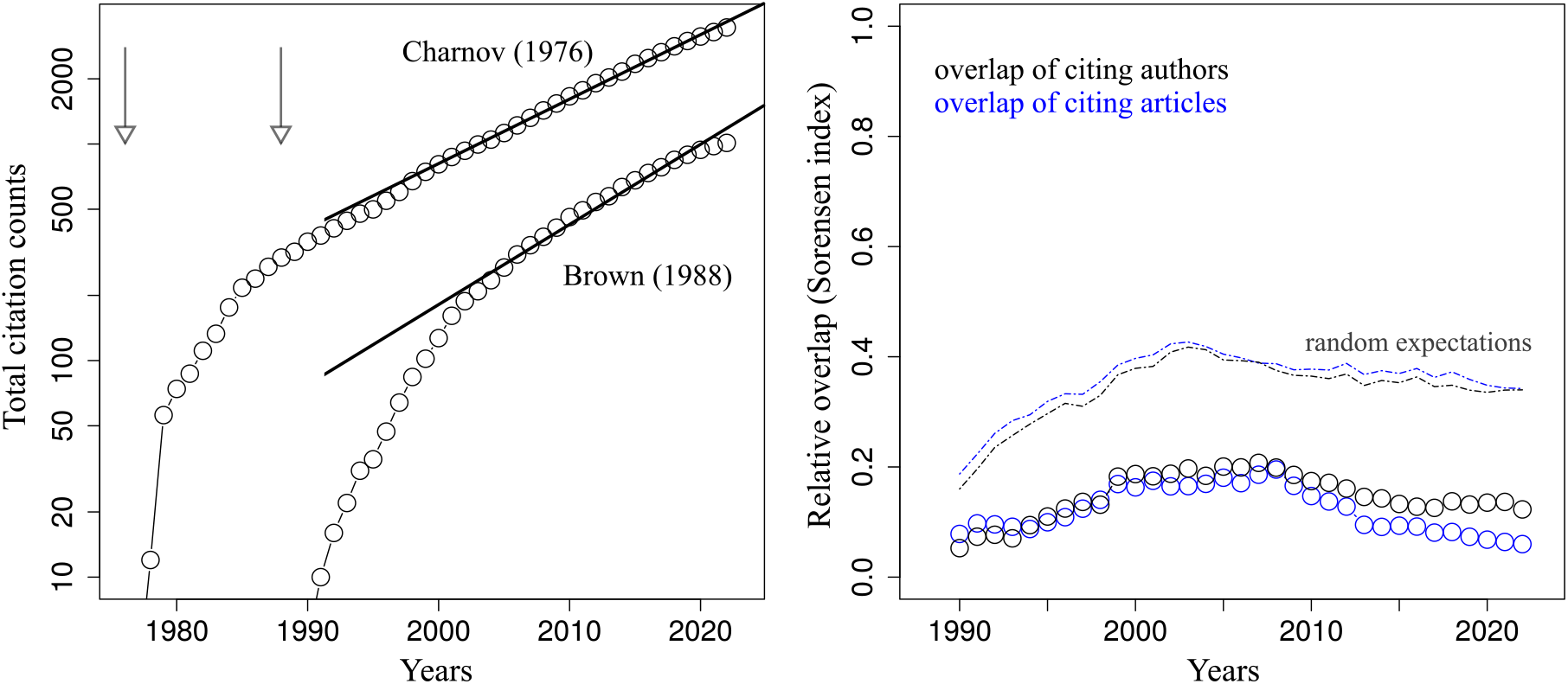
Charnov (1976) and Brown (1988): bibliometric facts. *Left*: Total number of citing articles as a function of time, from 1976. Note the logarithmic scale and the approximately linear (i.e. exponential) growth of citation counts in recent decades. Regression lines over the past 20 years are also shown. *Right*: Overlap (Sorensen index) in citing articles (black) and in citing authors (including all co-authors of citing articles; blue) from 1990. Indices are computed over sliding windows of five years. Dotted lines represent the values expected under chance, i.e. assuming independent citations.

We seek here to unify these two sister theories of patch-leaving decisions, by introducing the concept of risk into the original MVT. We first show that the MVT can readily accommodate fear effects and mortality risk, while retaining much of its predictive power and graphical appeal, proposing what we call the risk-MVT (rMVT). We then formally highlight the connections between this rMVT and Brown’s theory. We conclude with some simple predictions that can be derived from the rMVT. Our purpose is not to comprehensively study the consequences of risk on rMVT predictions, but to suggest a way to have the “resource acquisition” and “fear” perspectives of patch-leaving decisions reconciled in a single framework.

## MODELS AND METHODS

### Fitness maximization, GUD and Charnov’s original MVT

An individual forages over a large number of patches, a patch is characterized by some gain function *F*, and it takes on average *T* units of time to move from a patch to the next. We seek to determine how long to stay in a patch in order to maximize individual fitness. Fitness, in this context, is defined as the total amount of gains acquired after a long sequence of patch visits. If there is no risk of death, and more generally if the total duration of the sequence of patch visits is independent of the individual patch-leaving decisions (Calcagno 2018), then maximizing fitness is equivalent to maximizing the long-term rate of gain 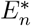 (Charnov 1976). *Here the subscript n* stands for “net gain”, emphasizing that energetic costs and losses should be discounted.

The MVT, in its general form allowing for heterogeneous patch characteristics (e.g. different resource levels; Calcagno et al. (2014)) defines the optimal residence times as:

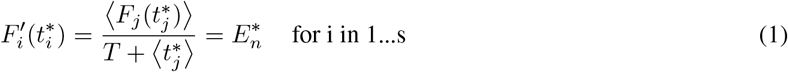

where *s* is the number of different patch types in the habitat, and *Fi* is the gain function in a patch of type *i*. Prime denotes differentiation with respect to time. We use brackets to denote averages over all patches, i.e. 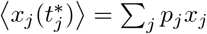, with *pj* the frequency of *j* patches, for any quantity *x* (Calcagno et al. 2013).

It follows from ((1)) the well-known result that patches should be left when the marginal rate of gain equals the long term average of gain in the habitat 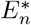. If all patches have identical characteristics, one can solve eq. ((1)) with a celebrated graphical solution involving a tangent line (Parker and Stuart 1976; Stephens and Krebs 1986; Calcagno 2018). Optimal residence times correspond to optimal levels of patch exploitation, as can be assessed by the amounts of resource left in patches, called giving-up densities or GUD (Brown 1988; Calcagno et al. 2014).

Eq. (1) is our “no-risk” starting point, in which we will seek to incorporate the effects of risk. In examples where all patches have identical characteristics, the *j* subscripts will often be dropped for clarity. In all examples, we will use the usual gain function *n*_0_(1 − exp (−*αt*)), with *n*_0_ the resource content of a patch and *α* the attack rate (Charnov 1976; Calcagno et al. 2013).

### The many facets of risk

Predation risk can impact foraging in many different ways (Brown et al. 1999; Arehart et al. 2023; Lima and Dill 1990). In the context of the MVT, three broad categories of risk ought to be distinguished, depending on the scale at which risk disrupts the foraging process. First, risk can impact solely the dynamics of resource acquisition at the local (within patch) level. For instance, the perception of risk can elicit vigilance and/or increase the level of stress while foraging in a patch (Apfelbach et al. 2005; Clinchy et al. 2013). We refer to these as “Disturbance” scenarios. Second, a more severe form of risk causes the premature end of patch visits, forcing the individual to leave the current patch. This applies to individuals that escape when perturbed by a potential predator, and to foragers that can be displaced by competitively dominant individuals or species (Blumstein 2006). We call these “Escape” scenarios. Last, risk can result in the death of the individual, an even more severe and global threat, as it prematurely stops not only the current patch visit but the entire foraging sequence. This puts “Death” scenarios in a league of their own. The three categories of risk have their impact on the MVT illustrated in Figure 2a, and are of course not mutually exclusive (Moll et al. 2017; Lima and Dill 1990).

**Figure 2.**
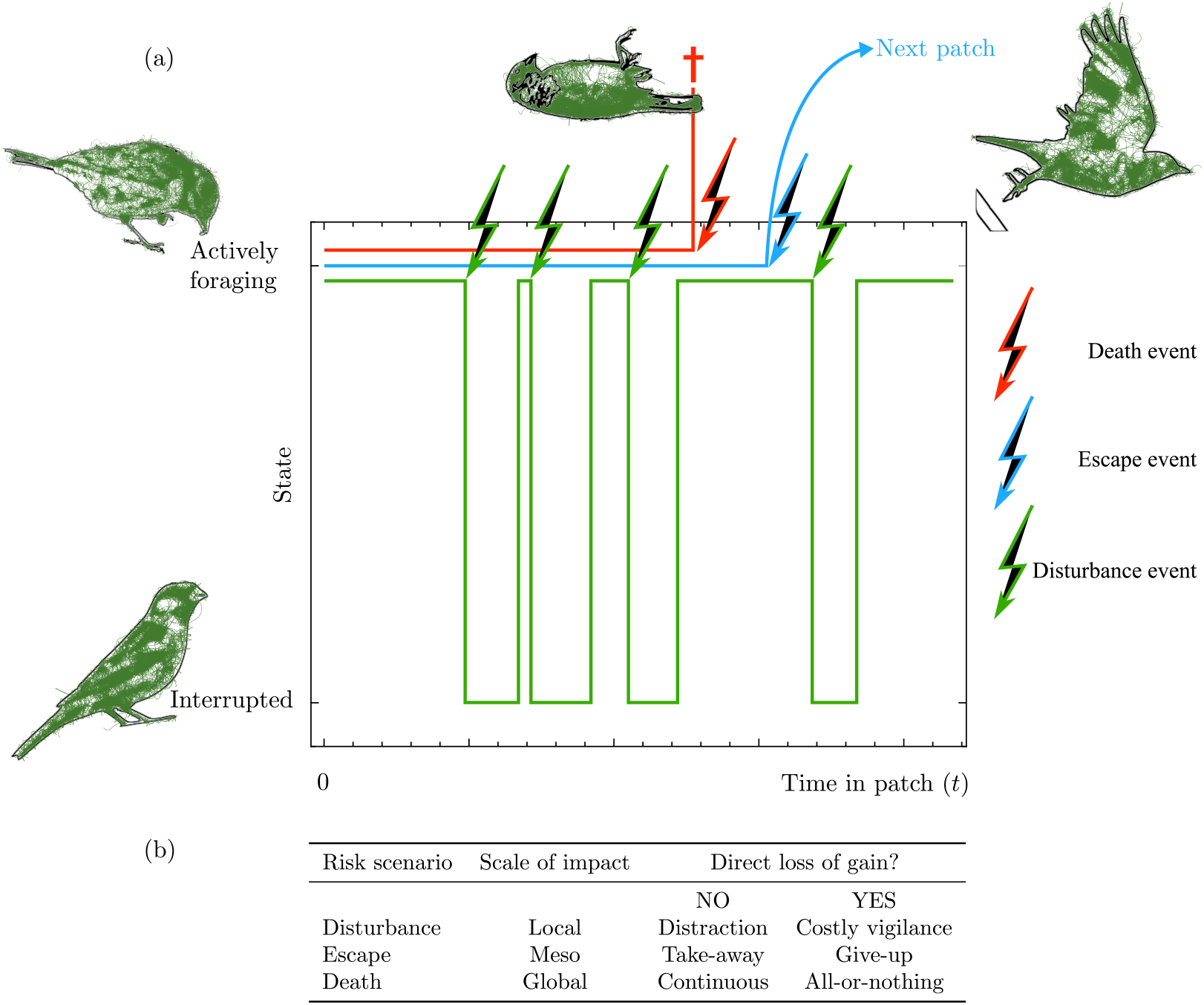
Introducing risk into the MVT. (a) Risk is modeled as stochastic events that disrupt the normal unfolding of a patch visit as time passes (x-axis). Three types of disruptions are possible. In Disturbance scenarios (green), each event (broken arrows) switches the individual from the actively foraging state to the interrupted state (y-axis). It takes some time for the individual to switch back to foraging. In Escape scenarios (blue), each event prematurely interrupts the patch visit and forces the individual to search for the next patch. Last, in Death scenarios (red), the event event kills the individual and interrupts both the current patch visit and the entire foraging sequence. (b) Depending on the type of risk, and on whether an interruption entails a direct loss of gains, we can identify six elementary risk scenarios. Of course all combinations and intermediate forms can occur in the real-world.

An additional distinction must be made, depending on whether the occurrence of risky events causes a direct loss of some, or even all, of the accumulated gains. Interruptions by predators, for instance, obviously reduce the effective foraging time, lowering future gain levels. This only interferes with the future dynamics of gain acquisition, an indirect loss. But dealing with predators may also imply an immediate expenditure of energy, or the loss of previously acquired food items, representing a direct loss. This is illustrated in Fig. 2b.

In this framework we can classify risk along two axes of severity: (i) the scale at which the normal course of events is disrupted, from local to global, and (ii) the amount of direct loss (Fig. 2b). This produces a huge spectrum of possibilities, but for clarity we will highlight six elemental scenarios, representative of familiar broadly-applicable situations. Disturbance scenarios can be of two extreme types: the transient interruption of foraging activities, without no direct loss (see Fig. 2b), will be called a pure “distraction” scenario. In contrast, risk can mostly have a diffuse direct cost (e.g. increased stress level), in what we call a “costly vigilance” scenario (Fig. 2b). In Escape scenarios, a natural contrast exists between escaping with its current patch harvest (“take-away” scenario; Fig. 2b) and escaping empty-handed (“give-up” scenario). Finally, in Death scenarios, an individual killed may retain all gains hitherto acquired, if gains are converted to fitness in a continuous manner, such as in parasitoids: all hosts parasitized by a female are potential offspring regardless of the female’s fate. We call this the “continuous” scenario (Fig. 2b), as opposed to the “all or nothing” scenario, where gains are valued in a delayed manner and only survivors benefit (see Arehart et al. 2023, for a similar classification).”All or nothing” is appropriate for a bee whose harvest is worthless if it does not make it back to the nest, or for species that must survive to the reproduction season if energy gains are to have any value.

In the following we consider how these different risk scenarios can be incorporated into the MVT. A Table of notations is provided in the Supp. Information (Table S1). Our general approach will be to (i) derive the expected (or realized) gain function, taking into account the effects of risk, (ii) determine the suitable fitness proxy, (iii) show what risk-MVT equations are appropriate to define optimal strategies. We will then show the connections between the risk-MVT and Brown’s GUD framework under the different risk scenarios. Finally we will present some predictions regarding the consequences of making a habitat more and more risky.

## INTRODUCING THE RISK-MVT

### Disturbance scenarios

We use a simple behavioral model, illustrated in Fig. 2a-b, according to which an individual in a patch can be in one of two states: (i) actively foraging, or (ii) engaged in some anti-predator activity (interrupted). The individual switches from the foraging state to the vigilant state at rate *β*_*j*_. Following an interruption, the individual reverts to the actively foraging state at rate *γ*_*j*_. This model can describe a range of particular behaviors, from scanning the neighbourhood when perceiving predator-related cues, to seeking protection in cover, dodging or deflecting attacks (Lima and Dill 1990; Werner et al. 1983). In all cases, 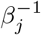 is the typical time interval to the next interruption and 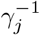 the typical delay to resume foraging following an interruption. In addition, each interruption may entail a one-off direct loss *δ*, representing for instance the energetic cost of performing a defensive behaviour.

The above model defines a two-state renewal process (Bénichou et al. 2005; Mangel 2006; Gallager 2013). From standard stochastic theory, we can compute the expected effective foraging time after some time *t* in a patch, 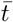, as well as its variance. We can from these express the expected gain of an individual spending time *t* in a patch of type *j* as:

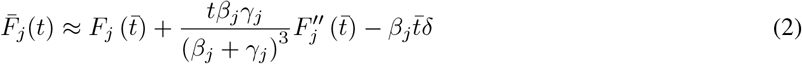

where *F*_*j*_ is the original gain function in the absence of risk (as in eq. (1)). This assumes *β*_*j*_ and *γ*_*j*_ are large enough, i.e. there are many relatively short interruptions in a typical patch visit (see Supp. Information for derivation).

Equation (2) describes a transformation of the original gain function, with three easy-to-interpret elements. The first term represents gains expected given the average time the individual is effectively foraging 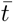. Up to a good approximation, the latter equals (*β* + *γ*(*β* + *γ*)*t*) */* (*β* + *γ*)^2^, omitting *j* subscripts for clarity (see S.I.). It accumulates slower than actual time, and therefore expected gains are less than *F*_*j*_(*t*), for the occurrence of interruptions. This effect is similar to that of decreasing the attack rate, or rate of resource consumption, commonly discussed in classical MVT theory (Charnov and Parker 1995). The second term is a variance effect: as the number of interruptions as well as their duration are random variables, so is the effective foraging time. The term 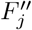 is typically negative, because the original gain function is typically concave (at least in the vicinity of optimal residence times; Charnov (1976)). Therefore stochasticity (variance in effective foraging time) will reduce the expected gain below 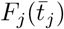, i.e. it introduces a cost of uncertainty. Finally, the third term in eq. (2) represents the direct cost of interruptions, proportional to the expected number of interruptions experienced and to *δ*, the cost per interruption. This third component is similar to adding an energetic cost of foraging (Stephens and Krebs 1986).

Eq. ((2)) can be simplified in the two special Disturbance scenarios highlighted in Fig. 2b. Under a pure “distraction” scenario, we set *δ* = 0, and provided *β*_*j*_ and *γ*_*j*_ are large enough, the second term can be neglected, leaving only the first. Conversely, we can recover the “costly vigilance” scenario (Fig. 2b) by setting *δ* to a low positive value and assuming *γ*_*j*_ is infinite, so that the individual pays a cost but does not pause foraging. The first term then simplifies to the original gain function *F*_*j*_(*t*), the second term vanishes, and the third boils down to −*βjδt*.

Provided one replaces the initial gain function *F*_*j*_ with the modified gain function defined by eq. (2), the MVT can be applied to characterize the optimal strategy. Examples are presented in Fig. 3a.

**Figure 3.**
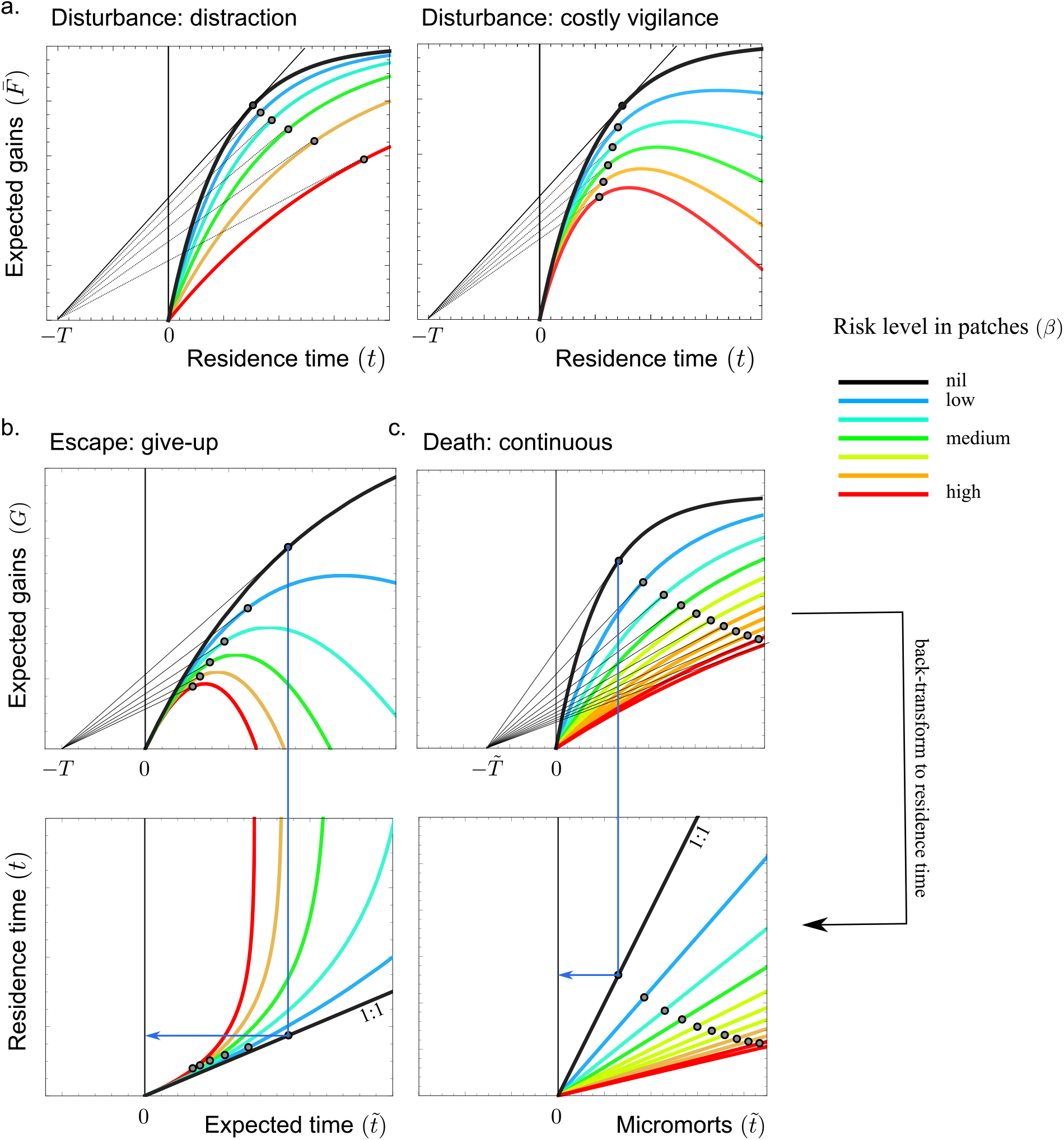
The risk-MVT. (a) In Disturbance scenarios, the optimization domain (x-axis) is regular time, and the gain function is modified to describe expected gains. (b) In Escape scenarios, the optimization domain is expected time 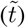, and the effective gain function is *G*, returning expected gains as a function of 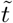. There is a one-to-one mapping between 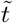 and regular time (bottom panel), so that one can back-transform one into the other. (c) In Death scenarios, 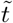is the amount of micromorts. In all cases, a homogeneous habitat is considered, with the initial gain function *F* and the level of risk *β* gradually increased from zero (blue to red). rMVT graphical solutions are represented (dark lines and circles), as well as the mapping with regular residence time in b and c (the mapping is just 1:1 in a).

### Escape scenarios

We now consider that an interruption does not simply suspend activity for a short while, but causes the individual to leave and search for another one (losing *T* units of time in the process; Fig. 2a). The individual may be forced to leave, or it may be optimal for it to leave (as if *γ*_*j*_ were very small in the previous scenario). Assuming as before that an interruption occurs at rate *β*_*j*_, two possibilities exist at time *t*: either the individual has not been troubled thus far, and has acquired *F*_*j*_(*t*) as in normal proceedings, or it was forced to leave at some time *ν < t*, so that it had acquired *F*_*j*_(*ν*) only, and paid a cost *δ*, if any. We can compute 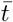, the expected foraging time of an individual that is willing to stay for time *t* in a patch. It takes value:

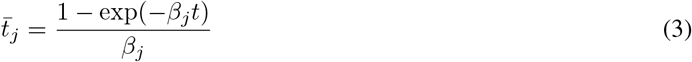

provided *β*_*j*_ is non-zero (see Supp. Information for derivation).

An individual willing to spend time *t* will stay for only 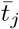 on average. Expected foraging time increases in a highly non-linear way with regular time *t* (eq. (3)). It plateaus at a maximum value of 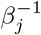 because in the long run interruption by a predator happens almost surely. The relationship between the two time-scales is illustrated in Fig. 3b. There is a one-to-one mapping between *t* and 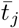, so by inverting equation (3) we can obtain a function 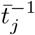 returning, for a given 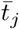, the corresponding *tj*.

Following the same approach as before, we compute the expected gains of an individual willing to stay in the patch for *t* time units as (see Supp. Information for derivation):

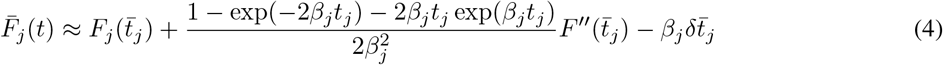

This again describes a transformation of the initial gain function, with the same three components as eq. (2). The first term represents a discounting effect: future gains are devalued because of the risk that an interruption occurs before. The second term represents the cost of uncertainty: as function *F* is typically concave, the gains effectively acquired on average, 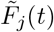, are less than 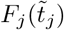, for the stochasticity in the time of interruption. The third term is the direct cost of an interruption, if any.

Setting *δ* = 0, i.e. dropping the third term, describes the “take-away” scenario in which an escaping individual retains all gains harvested in the patch. If, alternatively, an escaping individual must abandon any gain on site (“give-up” scenario; Fig. 2b), we set the direct cost *δ* to be the amount of gain acquired at the time of interruption *ν*, and the transformed gain function (4) simplifies to (see Supp. Information) *F*_*j*_(*t*) exp(−*β*_*j*_*t*). In that case, risk imposes a classical form of exponential discounting on the gain function.

A major difference from Disturbance scenarios is however that risk not only changes the expected gain function, but also changes the expected duration of patch visits. As a consequence, the long term average rate of gain 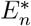 is still an appropriate fitness proxy, but is equal to

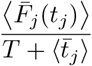

with expected residence times at the denominator.

An individual suffering from interruptions extracts lower gains from each patch visit (numerator), but also makes shorter patch visits on average (denominator). Fitness-wise, the latter effect partly counterbalances the former. Since the expression of 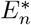 differs from that in the classical MVT (eq. (1)) we cannot use the expected gain function 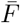 (eq. (4)) and simply plug it into the MVT equation (1), as we did for Disturbance scenarios.

We must further compute the gain function *G*, defined as

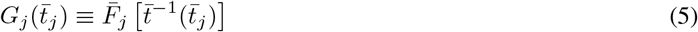

that returns the expected gain level, but as a function of the expected time 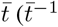 is obtained by inverting eq. (3)), rather than regular time *t*.

From this, we can rewrite the long term rate of gain as 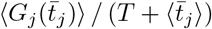, which depends on 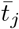 only. Optimizing this quantity is identical to maximizing 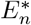 in the classical MVT, and so the optimal strategy can be determined using eq. (1). However the optimization domain is no longer *t*_*j*_ but 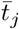 (eq. (3)) and *F* is replaced with *G* (eq. (5)). We shall call this a risk-MVT (or rMVT), as it generalizes the original equation on a different optimization domain. For generality, we will call the optimization domain 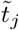; in this case 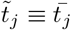, whereas we had 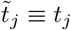 in Disturbance scenarios (as in the original MVT).

Up to this change, Escape scenarios are amenable to all usual MVT analyses, including graphical solutions and constructs. An example is provided in Fig. 3b.

### Death scenarios

We finally consider lethal interruptions, so that interruption events (still occurring at rate *β*_*j*_) now entail death, with no possibility to escape (Fig. 2a). In this case, we must also specify some background mortality rate *β*_*T*_, i.e. the risk of death when traveling between patches (Arehart et al. 2023). Over an entire sequence of patch visits, the average mortality rate is thus the arithmetic mean of instantaneous death rates, weighted by the fraction of time spent in each state:

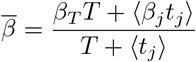

Unlike previous scenarios, predation risk here has no direct impact on the gain functions or the realized duration of patch visits, as it occurs by definition only once, and is a rare event at the scale of a typical patch visit. However, it affects the duration of the entire foraging sequence (*t*_*R*_), so that the long-term rate of gain 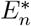 is no longer a suitable fitness proxy. An appropriate proxy is the expected amount of gain harvested in total over the entire sequence of patch visits (Brown 1988).

As long as there is no long-term trend in habitat quality, an individual acquires gain at an average rate of ⟨*F*_*j*_(*t*_*j*_) ⟩ */*(*T* + ⟨*tj* ⟩) until the end of the foraging sequence, risking death at rate 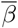. From this, the probability to survive until *t*_*R*_ is 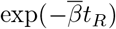, and if *t*_*R*_ is sufficiently large, the average lifespan is 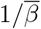. Obviously we must have *t*_*R*_ and 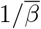 be much larger than *T* +⟨ *t*_*j*_⟩ to ensure that many patches are visited.

Maximizing the expected total gain is still quite remote from MVT-like maximization of 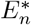, but an equivalence can be exhibited. First, in the “continuous” scenario, we let *t*_*R*_ go to infinity and assume that the gains acquired prior to death are all retained. Therefore the expected total gain reduces to 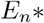 times the average life-span:

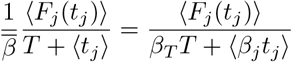

This is close to the MVT-like expression in eq. (1), except at the denominator we do not have regular times, but instead times weighted by the instantaneous level of risk (*β*_*j*_*t*_*j*_). Using the same approach as for Escape scenarios, we can change the optimization domain to the transformed scale 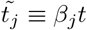 and 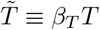. We then compute the corresponding transformed gain function 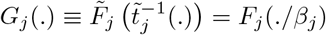. From this, we can determine the optimal strategy just as above.

If gains are converted to fitness only if death does not occur before the end of the entire foraging sequence (“all-or-nothing” scenario; Fig. 2c), it can be shown (see SI section 2.3) that the same rMVT applies, except that the hazard rate must be supplemented by 1*/t*_*R*_, representing the maximum duration of the entire foraging sequence (which is absent in the “continuous scenario”). In this case the result is only approximate, valid if the probability of surviving to the end of the foraging sequence is large enough (see SI for derivation).

The optimization domain suitable for Death scenarios thus differs from the one in Escape scenarios: 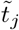 is a linear rescaling of time, with coefficient *β*_*j*_, as illustrated in Fig. 3c. It is not an expected duration in this case, and not even a time strictly speaking (it is dimensionless). In riskier patches, time is over-weighted as though risk accelerated the flow of time perceived by the individual (Fig. 3c). More formally, 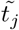 (resp. 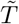) quantifies the infinitesimal increase in the probability of dying associated with a patch visit (resp. with traveling between patches). Such risk-penalized time, or dose of risk, is a common concept in the theory of decision analysis, known as the micromort unit introduced by Howard (Howard 1989, 2007). It is used to evaluate the relative dangerosity, defined as an increase in the probability of dying, of different human activities. In Death scenarios, the proper domain for the rMVT is thus the micromort scale, and optimal foragers can be seen as optimally allocating units of micromorts (rather than time units).

Death scenarios thus require to use micromorts as the rMVT optimization domain: scales 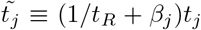 and 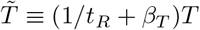. An example is provided in Fig. 3c.

### The risk-MVT in a nutshell

To summarize, all the above forms of risk can be handled as a risk-MVT (rMVT) formulation which generalizes the original MVT, with the same assumptions and formalism. The general equation for a rMVT is

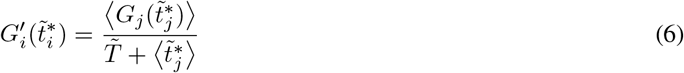

Here 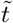is the effective optimization domain (the domain over which MVT-like rate maximization takes place). It may or may not be the regular time *t*, depending on the type of risk (see Table 1). *G* is the effective gain function, returning the expected gains as a function of 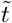. It can be obtained from a regular gain function in two steps. First, by modifying it to take into account the stochastic occurrence of risky events (interruptions, predation…). This step produced functions 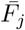 in the above above. Second, by expressing it on the effective domain 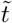, if different from the regular time *t*. One can then obtain the optimal residence time and optimal GUD by back-transforming 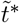 (Fig. 3). The original MVT (eq. (1)) is the special case where 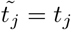 and *G*_*j*_ = *F*_*j*_.

**Table 1.** A summary of results. For the different forms of risk (rows), we indicated whether the problem is amenable to a risk-MVT formulation (“Charnovian”); the optimization domain on which the rMVT is to be formulated, and whether the effect of risk conforms to a predation cost as in Brown’s GUD framework (eq. (7); “Brownian”).

| Risk scenario | Charnovian? | rMVT domain* | Brownian? |
| --- | --- | --- | --- |
| None (original MVT) | YES | regular time | YES |
| Disturbance (distraction) | YES | regular time | NO |
| Disturbance (costly vigilance) | YES | regular time | YES |
| Escape (take-away) | YES | expected time | NO |
| Escape (give-up) | YES | expected time | NO |
| Death (continuous) | YES | micromorts | YES |
| Death (all or nothing) | YES<sup>†</sup> | micromorts | YES |

\*in full generality, the optimization domain is always "expected micromorts".
<sup>†</sup>good approximation if survival is high enough.

For clarity we have treated each risk scenario individually, but nothing prevents from applying the rMVT to a mixture of all types of risk (Moll et al. 2017; Lima and Dill 1990). For instance, if disturbance occurs at rate *β*_1*j*_, interruptions causing the individual to flee from the patch (Escape scenario) at rate *β*_2*j*_, and lethal predation (Death scenario) at rate *β*_3*j*_ (*β*_3*T*_ while traveling), one can describe the optimal strategy with the following rMVT. First, the optimization domain should be the product of expected residence time (as in Escape scenarios) and risk of death (as in Death scenarios). Therefore, one should use 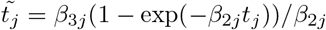, together with 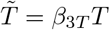. Those are “expected micromorts”.

The effective gain function *G*_*j*_ should be eq. (2) with *β*_1*j*_, to incorporate the effects of disturbance, then transformed according to eq. (3) with *β*_2*j*_, to account for escape effects, and finally transformed so as to be expressed as a function of the optimization domain 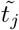 defined just above. Resulting equations may be tedious in those complex cases, but can be handled numerically or with usual mathematical tools. Note that the transformations involved are concavity-preserving, so that if the MVT can initially be applied with gain functions *F*_*j*_, the rMVT can as well (see S.I. Section 4).

### Connections to Brown’s GUD framework

In Brown’s GUD theory, an individual balances four different marginal rates when leaving a patch, so that the following equation holds

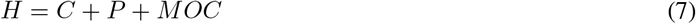

where *H* is the marginal rate of resource harvesting, *C* the marginal metabolic cost of foraging, *P* the cost of predation and *MOC* the “missed-opportunity cost”, i.e. a cost of not doing something else (Brown 1988). The equation is typically used within a heterogeneous habitat, characterized by some unknown (but fixed) *MOC*, to predict the consequences of varying parameter *P*. The most popular prediction is: in a riskier patch (*P* increases), the instantaneous rate of gain must be higher (*H* must increase), which implies a lower giving-up density (GUD) and a lower residence time (Brown 1988; Calcagno et al. 2014).

The first two quantities in eq. ((7)) are also involved in the MVT. The marginal rate of gain 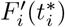 in eq. (1) is indeed a net gain, equal to the gross rate of energy gain minus the per-time unit energetic cost of foraging (Stephens and Krebs 1986), so that it equals *H* − *C*. The other two terms, *P* and *MOC* are lacking from the MVT, for obvious reasons. First, the MVT assumes there is no risk, *P* = 0. Second, as Brown’s equation focuses on a particular point in time with no particular global context, the *MOC* term is needed to justify that the individual is not doing something else at that time, and to ensure that the sum of marginal rates cancels out.

We can thus rewrite the classical MVT equation (1) as

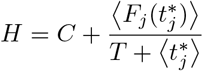

to exhibit the analogy with Brown’s equation.

Here the predation cost *P* is obviously zero, and the long-term average rate of gain (Charnov’s 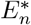; see eq. 1) can be seen to play the role of the *MOC*. Indeed, since the MVT provides a global model of patch visits, the *MOC* can be computed explicitly in terms of the gain functions, travel time, and optimal residence times. The alternative opportunity for the individual is “going to forage over some other patches”, which provides a rate of fitness gain equal to 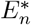, by definition.

Now the question is whether generalizing the MVT to predation risk introduces a fourth marginal rate: a predation cost *P >* 0 as in eq. (7). Obviously, under the costly vigilance scenario, we saw that risk introduced an additional cost term in the gain function, and it is indeed straightforward to formulate the MVT under “Brownian” form as

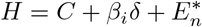

In the Death scenarios, the fitness proxy was the product of 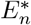, the long-term rate of gain, and a second term representing survival (either the average life-span or the probability of survival, depending on the scenario). If, as before, we split the net gain *F* into a gross harvest rate (*H*) minus a foraging cost (*C*), we can obtain the following expression (see SI section 3.1 for derivation):

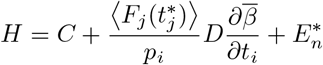

with *pi* the relative frequency of *i* patches and *D* the average duration of the entire foraging sequence, i.e. 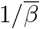 and *tR* in the continuous and all-or-nothing scenarios, respectively.

This again exhibits a full analogy with the “Brownian” form (eq. (7)), the *P* term involving the marginal change in the average death rate. Incidentally, the all-or-nothing scenario was the one assumed by Brown (1988), so it is expected to recover the same equation in that case.

Alas, in the other cases (distraction, including all Escape scenarios; Fig. 2c), the impacts of risk are more intricate and a neat four-fold additive partitioning cannot be achieved (see SI sections 3.2 and 3.3). This is because, even if we fix 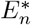, i.e. the *MOC*, increasing the level of risk not only impacts the predation cost *P*, but also the term *H* (and possibly, *C*) in eq. ((7)). This makes it impossible to vary parameter *P* without also varying the others, and the equation loses its applicability. These forms of risk are still “Charnovian” (rMVT compatible) but cannot be put in “Brownian” form. Forms of risks that are “Brownian” are those for which an increase in the level of risk has an additive effect on the instantaneous rate of gain. This can be because: (i) the effect is intrinsically additive, such as in the “costly vigilance” scenario, or (ii) the risk is sufficiently small at the level of one patch visit, leaving the expected gain functions essentially unchanged. The latter is what occurs in the “Death” scenarios.

The above results are summarized in Table 1.

### SOME RISK-MVT PREDICTIONS

We here highlight three results representative of the rMVT. We focus on a homogeneous habitat with all patches identical, and on the consequences of increasing the risk level in those patches. We will thus often omit *j* subscripts, noting for instance *β* for the level of risk in patches.

### Greater risk can have various effects on residence time and GUD

Greater risk in patches is expected to shorten optimal residence times, in a form of protection against risk (risk avoidance). Less directly, greater risk lowers the overall quality of the habitat, which from classical MVT theory increases residence time (Stephens and Krebs 1986; Calcagno et al. 2013). In the parlance of Brown’s GUD theory, increasing risk changes both *P* and *MOC*. It is therefore not straightforward to predict what the net effect should be (McNamara and Houston 1987). As shown in Fig. 4, increasing the level of risk in patches would very often decrease optimal residence time and increase optimal GUD, which is in accordance with the first intuition.

**Figure 4.**
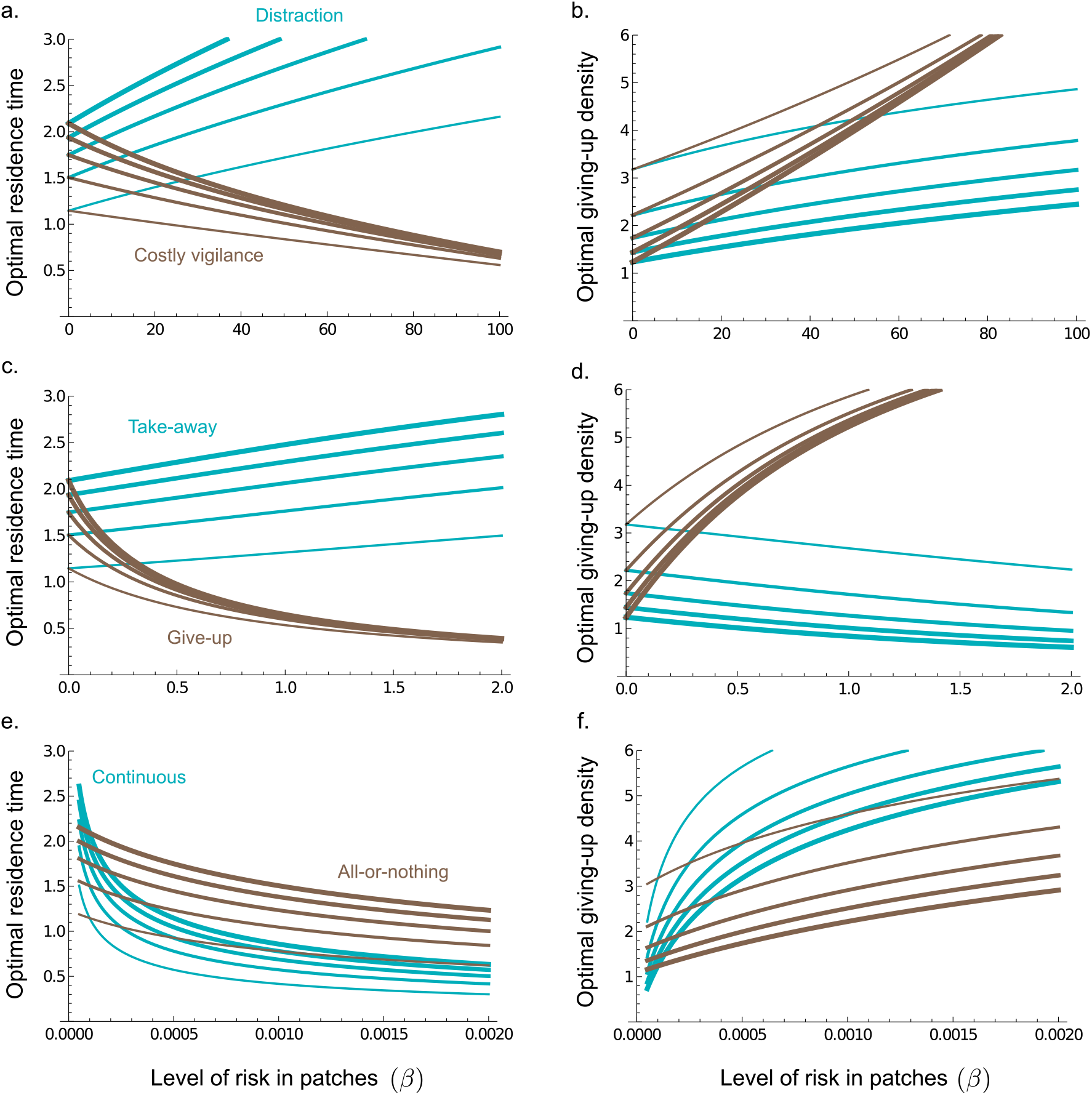
The impact of risk on optimal strategies. For every risk scenario, the rMVT optimal strategies is shown as a function of the level of risk *β* (x-axes), in terms of the optimal residence time *t*^*^ (left panels) or the optimal giving-up density (right panels). (a-b): Disturbance scenarios; (c-d): Escape scenarios; (d-e): Death scenarios. For each scenario, five values of the travel time *T* are used, ranging from 1 to 5 (increasing curve thickness). Other parameters: *γ* = 50 and *δ* = 0 (distraction), *γ* = 1000 and *δ* = 0.05 (costly vigilance), *β*_*T*_ = 1*e*^−4^ (Death scenarios), and *t*_*R*_ = 2000 (all-or-nothing). In all cases, *n*_0_ = 10 and *α* = 1.

There are certain exceptions however. In the Disturbance “distraction” and Escape “take-away” scenarios (in other words when predation risk mostly causes a loss of foraging time rather than an energetic loss), optimal residence typically increases with risk (Fig. 4a & c). Also, in the take-away scenario, optimal GUD decreases with risk (Fig. 4d). This reveals, in Disturbance scenarios, a decoupling of optimal residence time and GUD: the expected GUD depends not on *t*^*^, but on the optimal effective foraging time 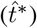, which can decrease with risk even though *t*^*^ increases (see Fig. S1). In such cases, although the optimal residence time *t*^*^ increases with risk, the optimal GUD increases as well, as it does in most other scenarios.

We remark that more severe forms of risk (e.g. Death versus Escape) require about one order of magnitude smaller risk intensity (*β*) to elicit comparable changes in the optimal strategies (compare the x-axes in Fig. 4). Some emblematic predictions of the MVT remain, reassuringly, valid in the presence of risk. For instance, optimal residence times are longer, and optimal GUDs lower, as travel time increases (Fig. 4).

### Predicted differences between experimental and field observations

The above results raise interesting implications for experimental studies of risk. A common type of experiments is indeed to observe the behavior of an individual in a context where predation is artificially suppressed, relative to the natural habitat (e.g. laboratory or field-enclosure experiments; Abrams (1990); Verdolin (2006)). In the rMVT, the optimal residence time *t*^*^ represents the residence time an individual is willing to adopt in a patch, but this does not necessarily represent the foraging time effectively realized: the occurrence of interruptions may shorten it. In other words, the optimal residence time (and the associated GUD) quantify the motivation, or persistence, of the individual, i.e. its intrinsic strategy, whereas foraging times observed in the field (and observed GUDs), quantify the practical outcomes of the strategy. The rMVT can be used to predict both.

As shown in Fig. 5, as the habitat gets riskier there can be a negative correlation between experimental observations (i.e. behaviours observed when predation risk is suppressed) and field observations (behaviours observed over several patches in the natural habitat). This occurs under the two risk scenarios for which optimal residence time increased with risk (distraction and take-away; see Fig. 4). In the take-away scenario for example (Fig. 5c & d), the individual should be more persistent when the habitat is riskier (*t*^*^ increases with *β*), but the average residence time observed in the field (where interruptions may occur; 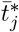, on the contrary decreases. The two measurements would thus differ, and the difference increases with the riskiness of the habitat (Fig. 5c). The same occurs if observing the effective foraging time 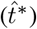 in the distraction scenario (Fig. 5a & b). Field observations would of course show some variability over different patches, but the predicted amount of variation suggests that differences would be statistically detectable, at least for high-enough levels of risk (Fig. 5).

**Figure 5.**
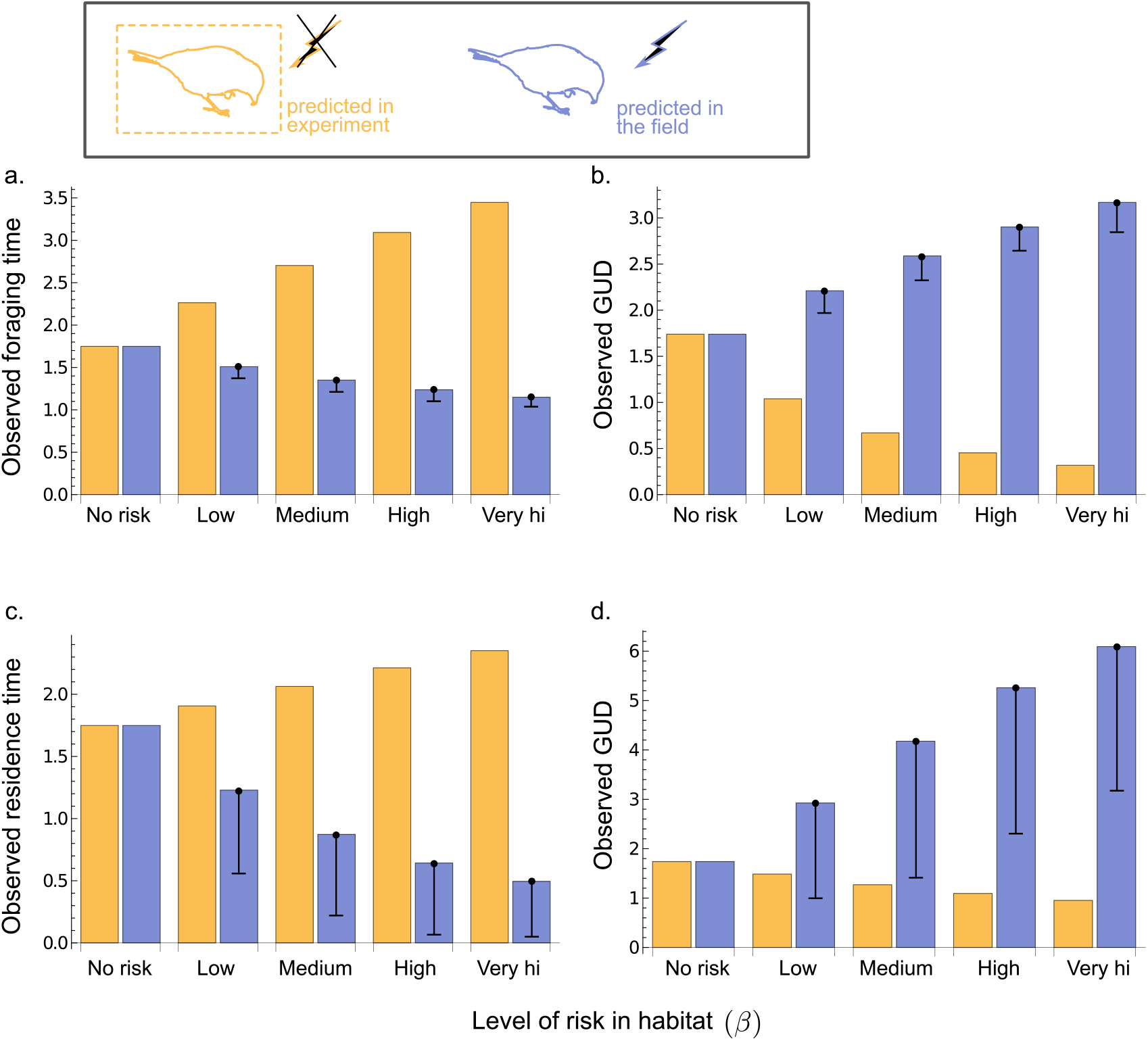
Experimental versus field predictions. In the Distraction (a-b) and Take-away (c-d) scenarios, rMVT optimality predicts contrasted patterns for experimental (orange) and field (blue) observations, as the level of risk in the habitat *β* increases (x-axis). Predictions are shown in terms of measurable times in the left panels (effective foraging time in a; residence time in c), and measurable GUD in the right panels. Experimental predictions apply to patch visits with experimentally suppressed predation risk, as explained in the main text. Field predictions apply to the expectation over several patch visits in the original habitat. Error bars represent one standard deviation, corresponding to different patch visits having different stochastic realizations of predator interruptions (experimental predictions have no such variability since risk is suppressed). *β* values range from 0 to 100 in (a-b), and from 0 to 2 in (c-d). Travel time is *T* = 3. Other parameters as in Fig. 4.

Another instance, in the Distraction scenario, is that an individual from a riskier habitat would stay longer and deplete the patch more importantly, when risk is artificially suppressed (Fig. 5a & b). This is because the absence of interruptions allows the individual to stay most of the time in the effectively foraging mode (see Fig. 2a), without losing time in the vigilant mode. Note that this prediction holds true if the leaving-rule mostly relies on evaluating elapsed time (time-based rule; Stephens and Krebs (1986); Ng et al. (2021)). If the individual rather mostly relies on evaluating resource density (GUD-based rule) or, equivalently, its effective foraging time (leaving when some prescribed foraging time is attained), the suppression of interruptions allows the target decision values to be reached more rapidly. Therefore in such experimental conditions the individual would leave earlier, and the differences between experimental and field observations shown in Fig. 5a-b would vanish. Other leaving rules might yield intermediate differences. This is an interesting situation where different patch-leaving mechanisms yield different testable predictions in terms of residence times, foraging times and GUDs.

### How daring individuals should be? Risk-avoidance and optimal boldness

Whilst micromorts quantify the dose of lethal risk in Death scenarios (Howard 1989), the rMVT naturally suggests extending them to all types of risk (lethal or not). We may compute the dose of risk taken as the optimal residence time *t*^*^ multiplied by the risk intensity *β*. In Disturbance scenarios, this quantifies the mean number of interruptions an individual suffers from per patch visit, and in Escape scenarios, the average number of interruptions an individual is in principle willing to endure. In all cases it is a quantitative measure of the amount of risk taken by an individual. Since the individual can adjust risk exposure by choosing the time spent in different patches (and which patches to ignore entirely), we regard this as a metric of how much risk it is willing to face, i.e. a metric correlated to boldness (Dammhahn and Almeling 2012; Harris et al. 2010; Wilson et al. 1994). We thus introduce “optimal boldness” as

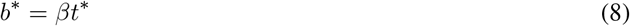

Remember that *t*^*^ is itself a (non-linear) function of *β*, so *b*^*^ does not simply convey the information contained in residence times. Of course, *b*^*^ coincides with 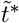 in Death scenarios. In heterogeneous habitats, the above definition extends directly as 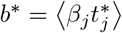, i.e. the mean dose of risk taken per patch visit.

Predictions for optimal boldness are shown in Figure 6. As expected, boldness is lower for more severe types of risk: it is maximal in Disturbance scenarios and minimal in Death scenarios, consistent with our classification of risk severity (Fig. 2b). Optimal boldness is also lower in risk scenarios entailing a direct loss of gain, or when travel time is shorter (i.e. in richer habitats). The latter result indicates that in habitats of lower quality, individuals are forced to take more risk in order to meet foraging requirements (Verdolin 2006).

**Figure 6.**
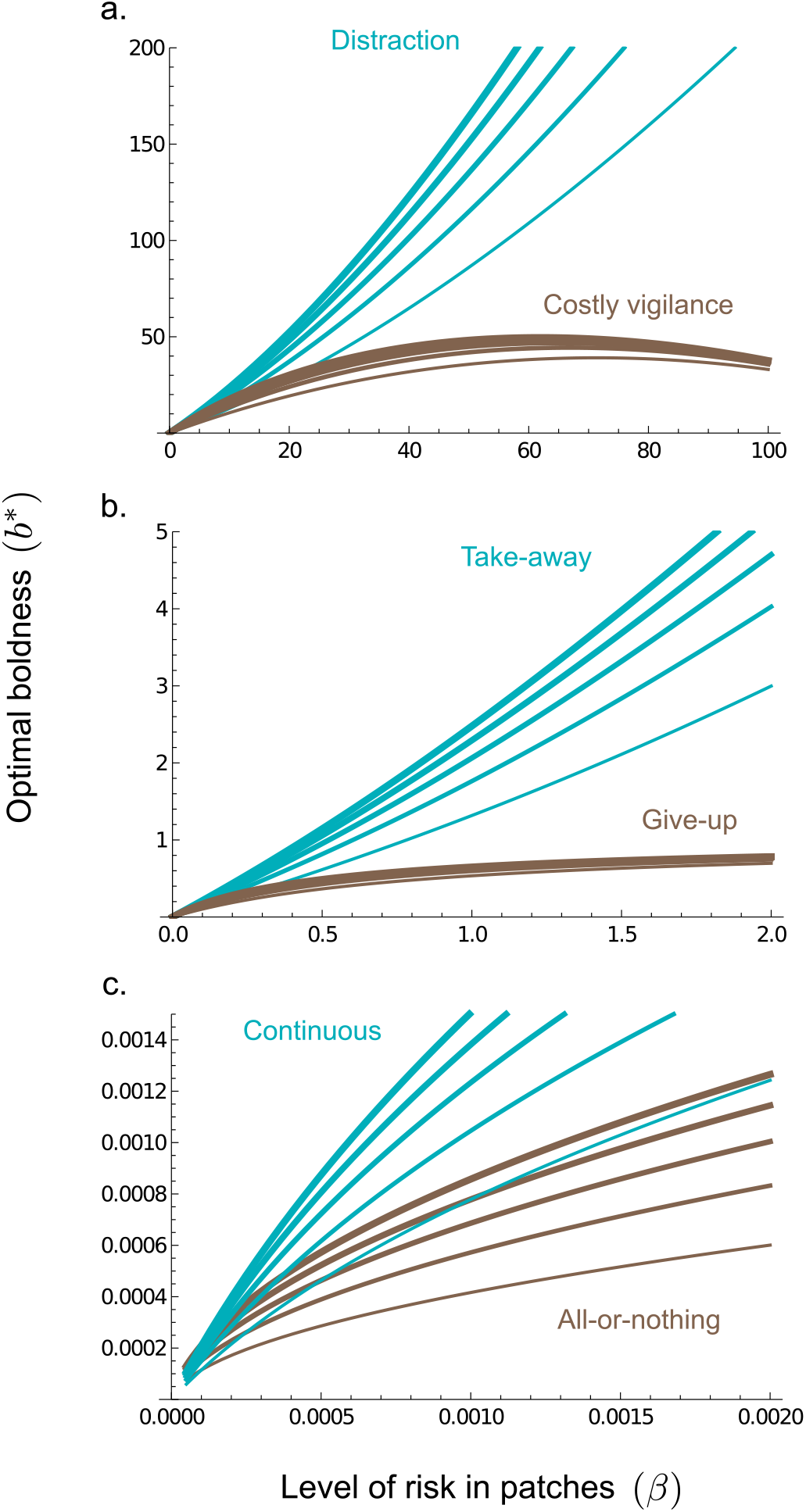
Optimal boldness. The optimal boldness *b*^*^, here quantified as the total amount of risk taken by an individual at the optimal strategy (*βt*^*^; eq. (8); y-axis) as a function of risk intensity *β*, for all risk scenarios. As previously, curves of increasing thickness correspond to travel time increasing from 1 to 5. Other parameters as in previous figures.

Most interestingly, optimal boldness very generally increases with the level of risk, across all scenarios. The only exception is when predation risk causes a direct loss of gain and is very intense, such as in the costly vigilance scenario with *β >* 60 (Fig. 6a). In such conditions, the consequences of risk become so dramatic that the individual shortens its residence time very importantly, in order to reduce risk exposure. In all other conditions, the rMVT predicts that an individual should accept to take more risks in a riskier habitat. If boldness did not vary with the level of risk, individuals would adopt more prudent strategies, with residence times scaling inversely with risk. This is not the case. Even though optimal strategies typically respond to risk by shortening residence time (Fig. 4), the response never fully compensates the increasing risk level: the net outcome is a greater overall exposure to risk, i.e. “bolder” individuals.

## CONCLUSION

Optimal foraging theory has produced two very successful frameworks to apprehend patch-leaving decisions. First, Charnov’s Marginal Value Theorem (MVT) elegantly captured the effect of patch quality and resource acquisition, but failed to consider predation risk. Subsequently, Brown’s GUD-theory introduced the notion of predation risk and risk avoidance, but did not retain all MVT ingredients. The end result is that the two theories have remained surprisingly disconnected, each used to address one or the other facet of the problem. As we have shown here, although there is a formal equivalence between the two theories, an important difference is that Brown’s theory is more abstract and does not specify a particular life-cycle or habitat context. In contrast, the MVT is derived from an explicit model of sequential patch visits, rendering the different components, as well as there covariations, computable. Brown’s equation thus includes the MVT as a special case, with the tremendous advantage of including predation risk, but at the price of reduced concreteness and predictive capacities. Even though more sophisticated and flexible frameworks exist that can describe optimal strategies with more realism (Tenhumberg et al. 2001; Davidson and El Hady 2019; Arehart et al. 2023), the conceptual simplicity of the MVT and of Brown’s deterministic theories remains unmatched. Since both have strengths and weaknesses, it would be desirable to combine them into a single framework.

In this article we have demonstrated that the MVT can be extended to make predictions under predation risk, producing what we call the risk-MVT (rMVT). The rMVT generalizes the MVT, retaining the same assumptions and methods. The concept of rate-maximization and associated techniques thus apply in the face of risk. The familiar graphical construct used to solve for optimal strategies still holds, just like for the original MVT (Fig. 3). And it remains true that, in a heterogenous habitats, all patches should be left when the marginal rate of gain equals some global value representative of habitat quality (eq. (6)). The main conceptual change when moving from MVT to rMVT is a change in the optimization domain, which needs not the regular residence time. Rather, some function of residence time that includes the intensity of risk 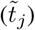 is the relevant scale against which rates of gain must be computed. One consequence is that time is not considered on equal footing in habitat portions with different levels of risk. This echoes the quite pervasive notion in biology that organisms do not weight time uniformly, but in a contextual manner. Examples include interval-timing in primates (Lake et al. 2016) or the degree-day timescale in ectotherm physiology (Charnov and Gillooly 2003). In full generality, we showed that the rMVT implies moving from residence time to expected dose of risk (formally micromorts; Howard (1989, 2007)) as the optimization domain: individuals are seen not as optimizing a rate of gain per unit of time, but per unit of mortality risk taken. The rMVT really predicts optimal micromort allocation, from which one can recover more usual metrics such as residence times, giving-up densities, expected foraging times, and others (Fig. 3). The intrinsic optimization domain of the rMVT (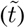) can always be regarded as expected micromorts, i.e. a dose of risk rather than a time. It just so happens that when death risk is absent (or invariant), one can simplify this domain down to expected time, or even regular time.

The rMVT also generalizes Brown’s GUD framework, in the sense that it is applicable to a broader range of risk scenarios. Death scenarios are the only ones for which Brown’s GUD theory is fully adequate (Table 1). Yet predators (or dominant competitors) can have a range of impacts on foraging individuals, beyond the obvious consumptive (mortality) effects (Moll et al. 2017; Arehart et al. 2023). Non-consumptive effects are pervasive, ranging from distraction to increased metabolic level to spatial relocation. Even though consumptive effects have historically received the greatest attention in ecology, non-consumptive effects are increasingly recognized are equally important, if not dominant (Peckarsky et al. 2008). This is especially true if one does not restrict its attention to predators, but includes competitors (from the same or other dominant species), as these seldom cause death but can still profoundly affect a forager’s behavior (Arseneau-Robar et al. 2022). The rMVT also brings an explicit modelling of the process of gain acquisition and the consequences of risk. The different marginal terms in GUD theory (eq. (7)) are intertwined and often cannot vary freely: their dependence on common underlying habitat characteristics necessarily introduces correlations between some of them. For instance, the *MOC* term is dependent on both *P* and *C*. When varying one habitat characteristic (such as the travel time, patch quality, foraging cost, or predation risk level) all four terms covary. The risk-MVT mechanistically describes the pattern of covariation and thus allows to predict the consequences of sustained changes in habitat parameters, such as the risk level *β*. In contrast, increasing the predation cost *P* in eq. (7), all else equal, cannot yield the kind of predictions presented here.

Predictions that can be derived from the risk-MVT exceed what this article could cover. One straightforward prediction is that a doubling of the death rate when searching for patches (*β*_*T*_) should, all else equal, have the same consequences as doubling travel time. This is because the effective travel time is simply *β*_*T*_ *T* (see “Death scenarios”). Another is that the consequences of increasing travel time, the result most emblematic of the MVT (Charnov 1976), remain unchanged in the presence of risk. It is likely that several predictions obtained under other frameworks could be rederived under the rMVT, and perhaps proven to hold under a broader range of risk scenarios. As the risk-MVT retains most of the MVT structure, it is amenable to all existing MVT methods, such as graphical constructs (Stephens and Krebs 1986), dimensional analysis (Stephens and Dunbar 1993; Charnov and Parker 1995) or sensitivity analysis (Calcagno et al. 2013). These would allow to investigate the impact of risk on optimal strategies, interactions with e.g. the distribution of resources or collective decision-making (Calcagno et al. 2014; Davis et al. 2022). For instance, the trade-offs between selecting resource rich versus safe patch could be investigated, or the effects of changes of prey handling time in response to risk (Verdolin 2006).

In this article we showed how different types of predation risk can have different consequences on optimal foraging strategies, as one might suspect (Moll et al. 2017; Arehart et al. 2023). The general expectation that greater risk in patches should select for shorter residence times (e.g. Dammhahn and Almeling (2012)) is often true, but the opposite can be optimal too, as is observed in Disturbance and Escape scenarios, if interruptions do not cause a direct loss of gain (Fig. 4). It follows that in general situations, with intermediate direct costs, one might observe non-monotonous trends in optimal residence time along gradients of risk, with an initial increase followed by a subsequent decrease, possibly looking like an absence of relationship (Fig. S2). Only in Death scenarios is a decrease of optimal residence time with risk always predicted. This suggests that in the (arguably more realistic) situations where several risk scenarios co-occur, different effects might cancel out and blur the association of risk level with optimal optimal residence times and GUDs.

An interesting set of predictions relates to the difference between the intrinsic persistence of individuals, as captured by the optimal residence time and GUD, and the actual realizations of patch visits in the presence of risk, as captured by the expected foraging times and expected GUDs. In some risk scenarios, greater risk selects for more persistent individuals, yet the expected residence time in natural condition would still diminish as risk increases (Fig. 5). The intrinsic persistence of individuals may be accessed experimentally, by artificially removing predation risk, and the actual realized foraging times and GUDs can be accessed from field measurements and observations. Both approaches are commonly employed (Verdolin 2006). The rMVT predicts the covariation, sometimes negative, of the two types of measurements along risk gradients, that could be tested using available data in several systems. Such comparison may yield insights into foraging strategies and the nature of risk.

Finally, the risk-MVT suggests to interpret optimal strategies in terms of risk-avoidance or “boldness” (Dammhahn and Almeling 2012; Wilson et al. 1994; Harris et al. 2010). We proposed to extend the micromort concept to all forms of predation risk, to obtain a quantitative metric of individual boldness, *b*^*^ defined as the quantity of risk an individual is willing to endure per patch visit ((8)). Variations in optimal boldness as risk increases indicate whether an individual under-compensates, accepting to endure more risk, or over-compensates, decreasing its overall exposure to risk. An interesting result is that under the rMVT, individuals generally under-compensate, and optimal strategies can thus be described as risk-prone, in the sense that optimal boldness increases with the level of risk and its severity (Fig. 6). Optimal boldness would only decline in some scenarios and for extreme risk levels. Maximizing resource intake thus appears to promote more boldness in risky habitats, which is consistent with other theoretical results (McNamara and Houston 1987), and with findings that individuals from risky habitats tend to be bolder (Harris et al. 2010).

The risk-MVT combines the strengths of Charnov’s classical MVT and Brown’s GUD theory. It opens a range of possible theoretical investigations and empirical evaluations. We suggest the rMVT can facilitate the integration of resource acquisition with risk avoidance, and become a valuable asset in the toolkit of optimal foraging theory.

## Acknowledgements

V.C. acknowledges funding from INRAE and Université Côte d’Azur (IDEX UCA-JEDI). The authors are grateful to E. L. Charnov for comments and suggested readings.

## Data accessibility

No new data was generated during this work. The R script and citation data used to generate the bibliometric Figure 1 are available at Zenodo (https://doi.org/10.5281/zenodo.10058431).

## Author contributions

V.C. conducted the research and wrote the manuscript. All coauthors were involved in discussing the ideas and in proof-reading the main manuscript. F.H. and L.M. reviewed the Supporting Information. Authors are listed in alphabetical order.

## Peer-review

The previous version of this preprint was peer-reviewed and recommended by PCI Evolutionary Biology. The present version brings in a slightly improved eq (2), as well as improved Supp. Information; everything else is unchanged. The peer-reviews and recommendation can be found at https://doi.org/10.24072/pci.evolbiol.100715.

## REFERENCES

Abrams PA. 1990. The effects of adaptive behavior on the type-2 functional response. Ecology:877–885.

Apfelbach R, Blanchard CD, Blanchard RJ, Hayes RA, McGregor IS. 2005. The effects of predator odors in mammalian prey species: a review of field and laboratory studies. Neuroscience & Biobehavioral Reviews. 29:1123–1144.

Arehart E, Reimer JR, Adler FR. 2023. Strategy maps: Generalised giving-up densities for optimal foraging. Ecology Letters. 26:398–410.

Arseneau-Robar TJM, Anderson KA, Vasey EN, Sicotte P, Teichroeb JA. 2022. Think fast!: Vervet monkeys assess the risk of being displaced by a dominant competitor when making foraging decisions. Frontiers in Ecology and Evolution. 10:775288.

Begon M L HJ, Townsend CR. 1996. Ecology: from individuals to ecosystems.

Bénichou O, Coppey M, Moreau M, Suet P, Voituriez R. 2005. Optimal search strategies for hidden targets. Physical review letters. 94:198101.

Blumstein DT. 2006. Developing an evolutionary ecology of fear: how life history and natural history traits affect disturbance tolerance in birds. Animal behaviour. 71:389–399.

Brown J. 1988. Patch use as an indicator of habitat preference, predation risk, and competition. Behav Ecol Sociobiol. 22:37–47.

Brown JS, Laundré JW, Gurung M. 1999. The ecology of fear: optimal foraging, game theory, and trophic interactions. Journal of Mammalogy. 80:385–399.

Calcagno V. 2018. The Marginal Value Theorem in a Nutshell. Fath, B.D. (Editor), Encyclopedia of Ecology, 2nd edition. Elsevier.

Calcagno V, Grognard F, Hamelin FM, Wajnberg É, Mailleret L. 2014. The functional response predicts the effect of resource distribution on the optimal movement rate of consumers. Ecology letters. 17:1570–1579.

Calcagno V, Grognard F, Wajnberg E, Mailleret L. 2013. How optimal foragers should respond to habitat changes: a reanalysis of the marginal value theorem. J Math Biol. 0:1–29.

Charnov E. 1976. Optimal foraging the marginal value theorem. Theor Popul Biol. 9:129–136.

Charnov E, Parker G. 1995. Dimensionless invariants from foraging theory’s marginal value theorem. Proc Natl Acad Sci USA. 92:1446–1450.

Charnov EL, Gillooly JF. 2003. Thermal time: body size, food quality and the 10 c rule. Evolutionary Ecology Research. 5:43–51.

Clinchy M, Sheriff MJ, Zanette LY. 2013. Predator-induced stress and the ecology of fear. Functional Ecology. 27:56–65.

Dammhahn M, Almeling L. 2012. Is risk taking during foraging a personality trait? a field test for cross-context consistency in boldness. Animal behaviour. 84:1131–1139.

Davidson JD, El Hady A. 2019. Foraging as an evidence accumulation process. PLoS computational biology. 15:e1007060.

Davis GH, Crofoot MC, Farine DR. 2022. Using optimal foraging theory to infer how groups make collective decisions. Trends in Ecology & Evolution. 37:942–952.

Gallager RG. 2013. Stochastic processes: theory for applications. Cambridge University Press.

Harris S, Ramnarine IW, Smith HG, Pettersson LB. 2010. Picking personalities apart: estimating the influence of predation, sex and body size on boldness in the guppy poecilia reticulata. Oikos. 119:1711–1718.

Howard RA. 1989. Microrisks for medical decision analysis. International Journal of Technology Assessment in Health Care. 5:357–370.

Howard RA. 2007. Advances in decision analysis, chapter 3: The foundations of decision analysis revisited.

Lake JI, LaBar KS, Meck WH. 2016. Emotional modulation of interval timing and time perception. Neuroscience & Biobehavioral Reviews. 64:403–420.

Lima SL, Dill LM. 1990. Behavioral decisions made under the risk of predation: a review and prospectus. Canadian journal of zoology. 68:619–640.

Mangel M. 2006. The theoretical biologist’s toolbox: quantitative methods for ecology and evolutionary biology. Cambridge University Press.

McNamara JM, Houston AI. 1987. Starvation and predation as factors limiting population size. Ecology. 68:1515–1519.

Moll RJ, Redilla KM, Mudumba T, Muneza AB, Gray SM, Abade L, Hayward MW, Millspaugh JJ, Montgomery RA. 2017. The many faces of fear: a synthesis of the methodological variation in characterizing predation risk. Journal of Animal Ecology.

Morris D, Davidson D. 2000. Optimally foraging mice match patch use with habitat differences in fitness. Ecology. 81:2061–2066.

Ng L, Garcia JE, Dyer AG, Stuart-Fox D. 2021. The ecological significance of time sense in animals. Biological Reviews. 96:526–540.

Pacheco-Cobos L, Winterhalder B, Cuatianquiz-Lima C, Rosetti MF, Hudson R, Ross CT. 2019. Nahua mushroom gatherers use area-restricted search strategies that conform to marginal value theorem predictions. Proceedings of the National Academy of Sciences. 116:10339–10347.

Parker G, Stuart R. 1976. Animal behavior as a strategy optimizer: evolution of resource assessment strategies and optimal emigration thresholds. Am Nat. 110:1055–1076.

Peckarsky BL, Abrams PA, Bolnick DI, Dill LM, Grabowski JH, Luttbeg B, Orrock JL, Peacor SD, Preisser EL, Schmitz OJ, et al. 2008. Revisiting the classics: considering nonconsumptive effects in textbook examples of predator–prey interactions. Ecology. 89:2416–2425.

Preisser EL, Bolnick DI, Benard MF. 2005. Scared to death? the effects of intimidation and consumption in predator– prey interactions. Ecology. 86:501–509.

Rosenberg NA. 2020. Fifty years of theoretical population biology. Theoretical Population Biology. 133:1–12.

Stephens D, Krebs J. 1986. Foraging theory. Princeton University Press.

Stephens DW, Dunbar S. 1993. Dimensional analysis in behavioral ecology. Behavioral Ecology. 4:172–183.

Tenhumberg B, Keller MA, Tyre AJ, Possingham HP. 2001. The effect of resource aggregation at different scales: optimal foraging behavior of cotesia rubecula. The American Naturalist. 158:505–518.

Verdolin JL. 2006. Meta-analysis of foraging and predation risk trade-offs in terrestrial systems. Behavioral Ecology and Sociobiology. 60:457–464.

Werner EE, Gilliam JF, Hall DJ, Mittelbach GG. 1983. An experimental test of the effects of predation risk on habitat use in fish. Ecology. 64:1540–1548.

Westneat D, Fox C. 2010. Evolutionary Behavioral Ecology. Oxford University Press, USA.

Wilson DS, Clark AB, Coleman K, Dearstyne T. 1994. Shyness and boldness in humans and other animals. Trends in Ecology & Evolution. 9:442–446.

