## Supplementary Information for "Taking fear back into the Marginal Value Theorem: the risk-MVT and optimal boldness"

#### 1. GENERAL ASSUMPTIONS

We follow the same assumptions as those underlying the MVT (Charnov 1976; Calcagno et al. 2013). The most important ones are:

- Every patch belongs to a certain class of patches, characterized by a gain function. The gain function can in principle have any shape, but is continuous, twice-differentiable and is increasing and concave for all times, or at least over part of $\mathbb{R}^+$.

- The individual is "omniscient" in the following sense: it knows the global characteristics of the habitat (travel time, gain functions, frequency of different classes of patches) and can determine the class of a patch when it enters one (and therefore the corresponding gain function).

- The individual can adjust its residence time in every class of patch, i.e. the time it will spend in a patch of a certain class, before leaving in search of the next patch.

- The individual will visit a long sequence of many patches, so that it experiences the statistical distribution of patches in the habitat, and seeks to maximize the total amount of gains acquired at the end of this long sequence. If the duration of the sequence is infinite, or finite but independent from the decisions made by the individual, then this is equivalent to maximizing the long-term expected rate of gain per unit of time.

Note that the first of the above assumptions would seldom hold strictly for a particular patch visit: although in some cases the process of resource consumption in a patch may be almost smooth and deterministic (e.g. a chimp sucking-up orange juice with a straw), in most real-world applications the process of resource acquisition during a particular patch visit will be discontinuous and stochastic. Most typically, a certain number of prey is to be captured in a patch, and the process of resource acquisition would occur as a sequence of steps, each step corresponding to the consumption of one prey. The exact times at which each consecutive prey captures take place would vary impredictably over different patch visits. The MVT only makes predictions for the optimal residence time in the *class of patches*, based on the expected average gain function of this class. The latter is indeed smooth and deterministic, and corresponds to the average over many particular stochastic realizations. Similarly, the rMVT makes predicions for a class of patches of a given risk level, and stochasticity in the occurence of specific interruptions by predators plays the same role as stochasticity in the times of prey capture in the previous example.

In all numerical illustrations and results, we will consider a homogeneous habitat in which every patch is characterized (in the absence of any risk) by the standard gain function:

$$F(t) = n_0 \left(1 - \exp(-\alpha t)\right)$$

This describes the consumption of $n_0$ prey items with some constant attack rate $\alpha$.

### 2. INCORPORATION OF RISK INTO THE MVT

#### 2.1. Disturbance scenarios

The two-state behavioral model illustrated in the main text (Fig. 1) is such that the individual is actively foraging at time $t$ with probability $p(t)$. This probability changes with time according to

$$p'(t) = \left(1 - p(t)\right)\gamma_j - p(t)\beta_j$$

prime indicating differentiation with respect to time.

In stochastic theory, this model defines an alternating Poisson process (Cox 1970; Gallager 2014), from which we can derive several quantities of interest.

As the individual enters the patch, it is initially in the actively foraging state, and so we solve from 0 to $t$ with initial condition $p(0) = 1$:

$$p(t) = \frac{\beta_j \exp\left(-(\beta_j + \gamma_j)t\right) + \gamma_j}{\beta_j + \gamma_j}$$

We are interested in the effective foraging time $\bar{t}$ (i.e. time spent in the foraging state), of an individual deciding to stay for time $t$ in a patch. To obtain the average foraging time, we can integrate as follows

$$\mathrm{E}(\bar{t}) = \int_0^t p(\tau) d\tau = \frac{\beta_j \left(1 - \exp\left(-t\left(\beta_j + \gamma_j\right)\right)\right) + t\gamma_j(\beta_j + \gamma_j)}{\left(\beta_j + \gamma_j\right)^2} \quad (1)$$

As soon as $t$ is long enough compared to the pace of interruptions, i.e. if the rates $\beta_j$ and $\gamma_j$ are high enough, we can obtain a pretty good approximation as

$$\mathrm{E}(\bar{t}) \approx \frac{\beta_j + t\gamma_j(\beta_j + \gamma_j)}{\left(\beta_j + \gamma_j\right)^2}$$

As time gets even larger, this further simplifies as

$$\mathrm{E}(\bar{t}) \approx \frac{t\gamma_j}{\beta_j + \gamma_j}$$

echoing the fact that, in the long run, the individual is actively foraging with probability $\gamma_j/(\beta_j + \gamma_j)$. The previous approximation is slightly better, for the initial condition, i.e. the fact that the individual is actively foraging when entering a patch, is better taken into account. For this reason we retained it.

Therefore, an individual deciding to spend time $t$ on the patch will on average forage for time $\mathrm{E}(\bar{t})$ only. For simplicity, we omitted the expectation symbol in the main text and used $\bar{t}$ directly. However, the actual foraging time is a random variable, and will vary for patch visit to patch visit. From standard renewal process theory (Cox 1970; eq. 16), we can further compute the variance of the effective foraging time as

$$\mathrm{var}(\bar{t}) = \frac{2t\beta_j\gamma_j}{(\beta_j + \gamma_j)^3} \tag{2}$$

which, if $\beta_j \approx \gamma_j$, can simplify to

$$\mathrm{var}(\bar{t}) \approx \frac{t}{4\beta_j}$$

Finally, the average number of times the individual is interrupted on the period is

$$\bar{n} = \frac{\beta_j\gamma_j t}{\beta_j + \gamma_j}$$

If every interruption has a one-off cost $\delta$, it follows that the direct loss of gain caused from interruptions is on average$\bar{n}\delta$. As before, these results assume that $t$ is large enough, but proved to have very good accuracy in all the cases considered.

From these results, we can approximate the effective gains acquired on average at time $t$ as

$$\bar{F}_j(t) = \mathrm{E}\left(F_j(\bar{t})\right) \approx F_j(\mathrm{E}(\bar{t})) + \frac{1}{2}F_j''\left(\mathrm{E}(\bar{t})\right)\mathrm{var}(\bar{t}) - \bar{n}\delta$$

following from a Taylor expansion at $\bar{t}$, up to second order.

This yields eq. (2) in the main article.

#### 2.2. Escape scenarios

If risk causes the individual to leave the current patch, we can proceed as above, by deriving a transformed gain function giving the gains effectively expected for an individual willing to stay in the patch for time $t$. Two possibilities exist at time $t$. First, it might not have been interrupted, and gain accumulation has proceeded as expected: it has acquired $F_j(t)$. Second, it may have been interrupted already, at some time $\nu < t$, in which case it has acquired gains equal to $F_j(\nu)$ only, rather than $F_j(t)$.

Assuming interruptions by predator occur at a constant rate $\beta_j$ the probability of having been interrupted by time $t$ is given by

$$1 - \exp(-\beta_j t)$$

We can thus express the gains effectively acquired on average by an individual that decides to stay for $t$ time units as

$$F_j(t)\exp(-\beta_j t) + \beta_j \int_0^t \exp(-\beta_j \nu) F_j(\nu) d\nu - (1 - \exp(-\beta_j t))\delta_j \quad (3)$$

This is the exact transformation describing the consequences of interruptions in Escape scenarios, but it is useful to derive an approximate form exhibiting the same structure as obtained previously in Disturbance scenarios.

The time at which the interruption occurs follows a truncated exponential distribution with parameter $\beta_j$. As for the Disturbance scenario, we can define $\bar{t}_j$, the expected time effectively spent foraging for an individual willing to stay for time $t$. It is the waiting time to the first interruption, and thus

$$\mathrm{E}(\bar{t}_j) = \beta_j \int_0^t \exp(-\beta_j \nu)\nu d\nu + t\exp(-\beta_j t) = \frac{1 - \exp(-\beta_j t)}{\beta_j}$$

The last simplification is eq. (3) in the main document and assumes risk is not entirely absent ($\beta_j > 0$).

We can further compute the variance of the effective residence time as

$$\mathrm{var}(\bar{t}_j) = \beta_j \int_0^t \exp(-\beta_j \nu)\nu^2 d\nu + t^2 \exp(-\beta_j t) - E(\bar{t}_j)^2$$

which can be expressed as

$$\mathrm{var}(\bar{t}_j) = \frac{1 - \exp(-2\beta_j t_j) - 2\beta_j t_j \exp(\beta_j t_j)}{2\beta_j^2}$$

Finally, knowing $E(\bar{t}_j)$ and $\mathrm{var}(\bar{t}_j)$, we can as before use a second-order Taylor expansion to approximate the average gain harvested at time $t$, yielding eq. (4) in the main document. The approximation is less accurate than in the Disturbance scenarios, because the underlying probability distribution is close to Gaussian in those scenarios, whereas it is closer to exponential in Escape scenarios. However we checked numerically that the approximation was satisfying. One can perfectly use the exact transformed function given above (eq. (3)) instead, with no further change, if favoring full accuracy over interpretability.

#### 2.3. Death scenarios

In Death scenarios, the fitness proxy to maximize is the total expected gain of an individual at its death, or at time $t_R$. Until either of these events, the individual is visiting patches and accumulating gains at rate $E_n^*$. Its expected gains are therefore

$$\frac{\langle F_j(t_j)\rangle}{T + \langle t_j\rangle} t$$

where $t$ is the relevant duration of the entire foraging sequence (depending on the Death scenario).

In the "continuous" scenario ($\delta = 0$ and $t_R \to \infty$), we get for the total expected gains:

$$\frac{1}{\overline{\beta}} \frac{\langle F_j(t_j)\rangle}{T + \langle t_j\rangle}$$

since $1/\overline{\beta}$ is the expected lifespan. From this, we get the rMVT derivation presented in the main document, involving the micromort scale as the effective optimzation domain.

If gains are converted to fitness only if the individual survives to the end of the foraging period $t_R$ ("all-or-nothing" scenario), the total expected gain is:

$$\frac{\langle F_j(t_j)\rangle\, t_R}{T + \langle t_j\rangle} \exp(-\overline{\beta} t_R)$$

reminding that $\exp(-\overline{\beta} t_R)$ is the survival probability.

The above very much resembles a regular MVT long term rate of gain, stated on the regular timescale, with all gain functions multiplied by the duration of the foraging session $t_R$ and the survival probability $\exp(-\overline{\beta} t_R)$. This seems a simple multiplicative transformation, simpler than the ones derived above, but there is a fundamental difference: since $\overline{\beta}$ is a function of all residence times, the gain function in one patch depends not only on the residence time in that patch and on travel time, but also on the residence times in all other patches. This makes the gain functions entangled, an important deviation from MVT theory. Therefore, in the "all-or-nothing" scenario, we can formulate the problem exactly as a rMVT in homogeneous habitats only, i.e. when all patches have similar characteristics. In such conditions, the rMVT can be stated on the regular time-scale, just as for Disturbance scenarios, using the transformed gain function:

$$\widetilde{F}(t) = F(t) \exp\left(-\overline{\beta} t_R\right)$$

However, if we assume that $\overline{\beta} t_R$ is small enough (i.e. if the probability of surviving is large enough, no smaller than about 75%), we can obtain a good approximation of the total expected gain as:

$$\left(\frac{1}{1/t_R + \overline{\beta}}\right) \frac{\langle F_j(t_j)\rangle}{T + \langle t_j\rangle}$$

We remark that this is very similar to the expression obtained for the "continuous" scenario, with the additional presence of the $1/t_R$ term at the denominator. The latter can be understood as the additional effect of the cap on the total duration of the sequence of patch visits (which cannot exceed $t_R$, whereas it is unbounded in the "continuous scenario"). Taking this into account, we may define the effective time-scale as

$$\tilde{t}_j = \left(\frac{1}{t_R} + \beta_j\right) t_j$$

and

$$\tilde{T} = \left(\frac{1}{t_R} + \beta_T\right) T$$

This is as in the previous scenario a micromort scale, but with the additional $1/t_R$ risk factor. We can thus proceed with the rMVT exactly as for the "continuous" scenario.

### 3. CONNECTION WITH BROWN'S FRAMEWORK

In this section we show how the risk-generalized MVT can, or cannot, be formulated under the Brownian form (eq. (7) in the main text). These results are summarized in Table 1 in the main document. Exceptionally, we start with the "all or nothing" Death scenario since it is the one originally assumed by Brown's GUD theory.

#### 3.1 Death scenarios

##### 3.1.1. All or nothing

We start with the "all or nothing" death scenario. Following the usual MVT derivation, we can determine the optimal residence times by setting all time-derivatives of the fitness proxy to zero. As explained in the main text, the appropriate fitness proxy in this case is the expected total gain. Writing the probability of surviving until the end of the entire foraging sequence as

$$S = \exp(-\bar{\beta} t_R),$$

we get for patches of class $i$:

$$\frac{\langle F_j(t_j) \rangle}{T + \langle t_j \rangle} \frac{\partial S}{\partial t_i} + S \frac{\partial}{\partial t_i} \left( \frac{\langle F_j(t_j) \rangle}{T + \langle t_j \rangle} \right) = 0 \tag{4}$$

The rightmost derivative is the quantity that is canceled under the original MVT. Here, it is not equal to zero and thus the rate of gain will not be maximized.

Dividing both sides by $S$:

$$\frac{\langle F_j(t_j) \rangle}{T + \langle t_j \rangle} \frac{\partial \log S}{\partial t_i} + \frac{\partial}{\partial t_i} \left( \frac{\langle F_j(t_j) \rangle}{T + \langle t_j \rangle} \right) = 0$$

Expanding the right most time-derivative:

$$\frac{\langle F_j(t_j) \rangle}{T + \langle t_j \rangle} \frac{\partial \log S}{\partial t_i} + p_i \frac{F_i'(t_i) \left(T + \langle t_j \rangle\right) - \langle F_j(t_j) \rangle}{\left(T + \langle t_j \rangle\right)^2} = 0$$

Note that to reduce clutter, the time derivative of $F$ was here expressed with the prime notation (as in the main text).

Multiplying by $T + \langle t_j \rangle$:

$$\langle F_j(t_j) \rangle \frac{\partial \log S}{\partial t_i} + p_i F_i'(t_i) - p_i \frac{\langle F_j(t_j) \rangle}{T + \langle t_j \rangle} = 0$$

Now we will exhibit the connection between this expression and Brown's GUD theory (eq. (7) in the main text). We first divide by $p_i$ and isolate the time derivative

$$F_i'(t_i) = -\frac{\langle F_j(t_j) \rangle}{p_i} \frac{\partial \log S}{\partial t_i} + \frac{\langle F_j(t_j) \rangle}{T + \langle t_j \rangle}$$

Decomposing $F'$ as gross gain (i.e. harvesting; noted $H$) minus foraging costs ($C$):

$$H_i(t_i) = C - \frac{\langle F_j(t_j) \rangle}{p_i} \frac{\partial \log S}{\partial t_i} + \frac{\langle F_j(t_j) \rangle}{T + \langle t_j \rangle}$$

This exhibits the connection with Brown's expression (7), and taking the log of $S$ yields the expression given in the main text.

Remark that using the expression for the probability of survival $S$ (see beginning of this section), we can be reformulate this as

$$\frac{1}{1 - t_R(\overline{\beta} - \beta_i)} F_i'(t_i) = \frac{\langle F_j(t_j) \rangle}{T + \langle t_j \rangle} \tag{5}$$

Equation (5) is close to a classical MVT equation, except for the first term on the l.h.s, representing the effect of risk. For all patches the r.h.s. is the average rate of gain, as in the original MVT, but the l.h.s. is not the quitting rate, rather a risk-penalized quitting rate. The first term is greater than one if

$$\beta_i < \overline{\beta}$$

i.e. if the focal patch presents lower risk than the average hazard rate.

##### 3.1.2. Continuous

We now follow exactly the same approach in the "continuous" scenario. Following the usual MVT derivation, we can determine the optimal residence times by setting all time-derivatives of the fitness proxy to zero (see above), which for patches of type $i$ yields

$$\frac{\langle F_j(t_j)\rangle}{T+\langle t_j\rangle}\frac{\partial}{\partial t_i}\left(\frac{1}{\overline{\beta}}\right)+\frac{1}{\overline{\beta}}\frac{\partial}{\partial t_i}\left(\frac{\langle F_j(t_j)\rangle}{T+\langle t_j\rangle}\right)=0 \tag{6}$$

The rightmost derivative is the quantity that is canceled under the original MVT. Here, it is not equal to zero and thus the rate of gain will not be maximized.

Multiplying both sides by $\overline{\beta}$:

$$\frac{\langle F_j(t_j)\rangle}{T+\langle t_j\rangle}\frac{\partial \log}{\partial t_i}\left(\frac{1}{\overline{\beta}}\right)+\frac{\partial}{\partial t_i}\left(\frac{\langle F_j(t_j)\rangle}{T+\langle t_j\rangle}\right)=0$$

Expanding the right most time-derivative:

$$\frac{\langle F_j(t_j)\rangle}{T+\langle t_j\rangle}\frac{\partial \log}{\partial t_i}\left(\frac{1}{\overline{\beta}}\right)+p_i\frac{F_i'(t_i)\,(T+\langle t_j\rangle)-\langle F_j(t_j)\rangle}{(T+\langle t_j\rangle)^2}=0$$

Multiplying by $T+\langle t_j\rangle$

$$\langle F_j(t_j)\rangle \frac{\partial \log}{\partial t_i}\left(\frac{1}{\bar{\bar{\beta}}}\right) + p_i F_i'(t_i) - p_i \frac{\langle F_j(t_j)\rangle}{T + \langle t_j\rangle} = 0$$

Dividing by $p_i$ and isolating time derivative

$$F_i'(t_i) = -\frac{\langle F_j(t_j)\rangle}{p_i}\frac{\partial \log}{\partial t_i}\left(\frac{1}{\bar{\bar{\beta}}}\right) + \frac{\langle F_j(t_j)\rangle}{T + \langle t_j\rangle}$$

Decomposing $F'$ as gross gain $H$ minus foraging costs $C$:

$$H_i(t_i) = C - \frac{\langle F_j(t_j)\rangle}{p_i}\frac{\partial \log}{\partial t_i}\left(\frac{1}{\bar{\bar{\beta}}}\right) + \frac{\langle F_j(t_j)\rangle}{T + \langle t_j\rangle}$$

This exhibits the connection with Brown's framework (eq. (7) in the main text).

Expanding the remaining time derivative and rearranging we can define the optimal strategy as

$$\frac{\langle F_j(t_j)\rangle}{T + \langle t_j\rangle} p_i \frac{\beta_T T + \langle \beta_j t_j\rangle - \beta_i\left(T + \langle t_j\rangle\right)}{\left(\beta_T T + \langle \beta_j t_j\rangle\right)^2} + \frac{T + \langle t_j\rangle}{\beta_T T + \langle \beta_j t_j\rangle}\left(p_i \frac{F_i'(t_i)\left(T + \langle t_j\rangle\right) - \langle F_j(t_j)\rangle}{\left(T + \langle t_j\rangle\right)^2}\right) = 0$$

$$E_n^* \frac{\beta_T T + \langle \beta_j t_j\rangle - \beta_i\left(T + \langle t_j\rangle\right)}{\left(\beta_T T + \langle \beta_j t_j\rangle\right)^2} + \frac{1}{\beta_T T + \langle \beta_j t_j\rangle}\frac{F_i'(t_i)\left(T + \langle t_j\rangle\right) - \langle F_j(t_j)\rangle}{\left(T + \langle t_j\rangle\right)} = 0$$

$$E_n^* \frac{\beta_T T + \langle \beta_j t_j\rangle - \beta_i\left(T + \langle t_j\rangle\right)}{\beta_T T + \langle \beta_j t_j\rangle} + F_i'(t_i) - E_n^* = 0$$

$$F_i'(t_i) = -E_n^* \frac{\beta_T T + \langle \beta_j t_j \rangle - \beta_i \left(T + \langle t_j \rangle\right)}{\beta_T T + \langle \beta_j t_j \rangle} + E_n^*$$

$$F_i'(t_i) = E_n^* \frac{\beta_i \left(T + \langle t_j \rangle\right)}{\beta_T T + \langle \beta_j t_j \rangle}$$

$$F_i'(t_i) = \frac{\beta_i \left\langle F_j(t_j) \right\rangle}{\beta_T T + \langle \beta_j t_j \rangle}$$

$$\frac{F_i'(t_i)}{\beta_i} = \frac{\left\langle F_j(t_j) \right\rangle}{\beta_T T + \langle \beta_j t_j \rangle}$$

In a heterogeneous habitat, all patches will have the same quitting rate on the effective time-scale (micromorts), which on the regular timescale implies that all patches will have the same value of the risk-penalized quitting rate:

$$F_i'(t_i)/\beta_i$$

This simple form of risk-penalization has been previously called the $\mu/f$ rule (Brown et al. 1992).

#### 3.2. Escape scenarios

We now apply the same approach to the Escape scenarios. Let us introduce quantity

$$A = \frac{T + \langle \bar{t}_j \rangle}{T + \langle t_j \rangle}$$

that represents the "rescaling" of time caused by risk, i.e. the factor by which the duration of one patch-cycle is shortened because of premature interruptions. Conversely, the inverse of $A$ represents the average number of patches that are visited in the presence of risk relative to in the absence of risk, over a given period of time. Hence we can re-express fitness as

$$\frac{1}{A}\frac{\left\langle \bar{F}_j(t_j)\right\rangle}{T+\langle t_j\rangle}$$

Following the usual MVT derivation, we can determine the optimal residence times by setting all time-derivatives of the fitness proxy to zero, which for one patch $i$ yields

$$\frac{\left\langle \bar{F}_j(t_j)\right\rangle}{T+\langle t_j\rangle}\frac{\partial}{\partial t_i}\left(\frac{1}{A}\right)+\frac{1}{A}\frac{\partial}{\partial t_i}\left(\frac{\left\langle \bar{F}_j(t_j)\right\rangle}{T+\langle t_j\rangle}\right)=0$$

Following the same calculations as before we get

$$\bar{F}_j'(t_i)=\frac{\left\langle \bar{F}_j(t_j)\right\rangle}{T+\langle t_j\rangle}-\frac{\left\langle \bar{F}_j(t_j)\right\rangle}{p_i}\frac{\partial \log}{\partial t_i}\left(\frac{1}{A}\right)$$

However, since the transformed gain functions are involved in the equation, we cannot partition from the l.h.s. the cost of foraging from the harvest rate. Furthermore, risk parameters are also present in the time-derivative, and cannot be disentangled from the others. Therefore we cannot reach a partition of terms *à la* Brown.

#### 3.3. Disturbance scenarios

It is immediately obvious that in the "costly vigilance" scenario, the additional energetic loss introduced by risk ($-\beta_j\delta$) takes the role of $P$ in main text eq. (7).

In the "distraction" scenario, the action of risk is to cause a linear rescaling of time. This does not have a simple additive effect on the gain function: it is equivalent to decreasing the rate of resource harvesting, and an additional $P$ term does not result. Since increasing the level of risk in a patch correlatively alters the shape of the harvest (and possibly foraging cost) functions. As above for the Escape scenarios, Therefore we cannot reach a partition of terms *à la* Brown.

### 4. ON THE CURVATURE OF $G$

Starting from a gain function $F$, meeting the usual requirements of the MVT (see Section 1), the rMVT implies to transform it into expected gains, and then, possibly, by restating the problem on the effective optimization domain. These transformations alter the concavity properties of the initial gain function $F$, and we must verify that a fitness maximum still exists, in order to be able a MVT-like equation stated in terms of the effective gain function $G$. Let us consider the two steps one after the other.

#### 4.1. Moving to expected gains

Let us first consider Disturbance scenarios. The inclusion of risk produces the expected gain function:

$$\bar{F}_j(t) = F_j\left(\frac{\gamma_j t}{\beta_j+\gamma_j}\right) + \frac{t\beta_j\gamma_j}{(\beta_j+\gamma_j)^3}F_j''\left(\frac{\gamma_j t}{\beta_j+\gamma_j}\right) - \frac{\beta_j\gamma_j\delta}{\beta_j+\gamma_j}t$$

Differentiating with respect to $t$ yields

$$\bar{F}_j'(t) = \frac{\gamma_j}{z_j^4}\left[z_j^3 F_j'\left(\tilde{t_j}\right) + \beta_j\left(z_j F_j''\left(\tilde{t_j}\right) + \gamma_j t F_j'''\left(\tilde{t_j}\right) - \delta z_j^3\right)\right]$$

with $\tilde{t_j} = \frac{\gamma_j t}{\beta_j+\gamma_j}$ and $z_j = \beta_j + \gamma_j$.

Therefore

$$\lim_{t\to\infty}\bar{F}_j'(t) \le 0$$

given that $\lim F_j'(t) \le 0$ (because patches can be exhausted), and under the mild assumption that $F_j''(t)$ and $F_j'''(t)$ both tend to zero.

Furthermore

$$\bar{F}_j'(0) > 0$$

as any patch worth exploiting must yield positive gains.

From continuity arguments, this implies that $\bar{F}_j(t)$ is increasing and concave over at least part of $(0,\infty)$, so that it also satisfies the concavity requirements of the MVT.

In Escape scenarios, the expected gains are, in their exact form:

$$\bar{F}_j(t) = F_j(t)\exp(-\beta_j t) + \beta_j \int_0^t \exp(-\beta_j \nu) F_j(\nu) d\nu - (1 - \exp(-\beta_j t))\delta_j$$

From this,

$$\bar{F}_j'(t) = \left(F_j'(t) - \beta_j \delta_j\right)\exp(-\beta_j t)$$

This indicates that if $F_j'(t)$ tends to a non-positive value, so does $\bar{F}_j'(t)$. Therefore, using the same argument as above, we can conclude that if $F_j(t)$ meets the concavity requirements, $\bar{F}_j(t)$ does too.

#### 4.2. Moving to the effective optimization domain $\tilde{t}$

The second change that may be involved in the rMVT is expressing $\bar{F}_j(t)$ as a function of the effective optimization domain $\tilde{t_j}$, yielding the effective gain function $G_j$:

$$G_j(\tilde{t_j}) \equiv \bar{F}_j\left(\tilde{t_j}^{-1}(\tilde{t_j})\right)$$

Function $G_j$ should be increasing and concave on at least part of its definition domain for the rMVT equation to be applicable.

We first remark that $\tilde{t_j}$ is a linear function of $t$ in Death scenarios, so the change from time to micromorts does not change the concavity properties: if $\bar{F}_j$ is concave, $G_j$ will be too.

This leaves us with Escape scenarios, for which $\tilde{t_j}$ is a concave increasing function of $t$:

$$\tilde{t_j} = \frac{1 - \exp(-\beta_j t)}{\beta_j}$$

The domain of definition of $G_j$ is

$$\left[0, \tilde{t_j}(\infty)\right) = \left(0, \frac{1}{\beta_j}\right)$$

From the definition of $G_j$ we have

$$G_j'(\tilde{t_j}) = \bar{F}_j'\left(\tilde{t_j}^{-1}(\tilde{t_j})\right)\left(\frac{d\tilde{t_j}^{-1}}{d\tilde{t_j}}\right)$$

As

$$\frac{d\tilde{t_j}^{-1}}{d\tilde{t_j}}(0) = 1$$

it follows that

$$G_j'(0) = \bar{F}_j'(0) > 0$$

Furthermore

$$\lim_{\tilde{t_j} \to 1/\beta_j} \left(\frac{d\tilde{t_j}^{-1}}{d\tilde{t_j}}\right) = \infty$$

and therefore

$$\lim_{\tilde{t_j} \to 1/\beta_j} G_j'(\tilde{t_j}) \leq 0$$

since $\lim \bar{F}_j'\left(\tilde{t_j}^{-1}(\tilde{t_j})\right) \leq 0$.

From the same continuity argument as before, this implies that $G_j$ is increasing and concave on at least part of its domain of definition, so that the rMVT can be applied.

### REFERENCES


- Brown, J. S., Morgan, R. A. & Dow, B. D. (1992) Patch use under predation risk: II. A test with fox squirrels, *Sciurus niger*. *Annales Zoologici Fennici* (pp. 311-318)

- Calcagno V., Mailleret, L., Wajnberg, É. & Grognard, F. (2014) How optimal foragers should respond to habitat changes: a reanalysis of the Marginal Value Theorem. *Journal of Mathematical Biology.* 69, (pp. 1237-1265).

- Charnov, E. L. (1976) Optimal foraging, the marginal value theorem. *Theoretical Population Biology*, 9(2), (pp. 129-136).

- Cox, D. R. (1970) *Renewal Theory.* Methuen & co, ltd, London.

- Gallager, R. G. (2014) *Stochastic Processes: Theory for Applications.* Cambridge University Press, Cambridge.
